# Enhancer Reprogramming in Melanoma Immune Checkpoint Therapy Resistance

**DOI:** 10.1101/2022.08.31.506051

**Authors:** Mayinuer Maitituoheti, Alvin Shi, Ming Tang, Li-Lun Ho, Christopher Terranova, Kyriaki Galani, Emily Z. Keung, Caitlin A. Creasy, Manrong Wu, Jiajia Chen, Nana Chen, Anand K. Singh, Apoorvi Chaudhri, Nazanin E. Anvar, Giuseppe Tarantino, Jiekun Yang, Sharmistha Sarkar, Shan Jiang, Jared Malke, Lauren Haydu, Elizabeth Burton, Michael A. Davies, Jeffrey E. Gershenwald, Patrick Hwu, Alexander Lazar, Jaime H. Cheah, Christian K. Soule, Stuart S. Levine, Chantale Bernatchez, Srinivas V. Saladi, David Liu, Jennifer Wargo, Genevieve M. Boland, Manolis Kellis, Kunal Rai

## Abstract

Immune checkpoint blockade (ICB) therapy has improved long-term survival for patients with advanced melanoma. However, there is critical need to identify potential biomarkers of response and actionable strategies to improve response rates. Through generation and analysis of 148 chromatin modification maps for 36 melanoma samples from patients treated with anti-PD- 1, we identified significant enrichment of active enhancer states in non-responders at baseline. Analysis of an independent cohort of 20 samples identified a set of 437 enhancers that predicted response to anti-PD-1 therapy (Area Under the Curve of 0.8417). The activated non-responder enhancers marked a group of key regulators of several pathways in melanoma cells (including c- MET, TGFβ, EMT and AKT) that are known to mediate resistance to ICB therapy and several checkpoint receptors in T cells. Epigenetic editing experiments implicated involvement of c-MET enhancers in the modulation of immune response. Finally, inhibition of enhancers and repression of these pathways using bromodomain inhibitors along with anti-PD-1 therapy significantly decreased melanoma tumor burden and increased T-cell infiltration. Together, these findings identify a potential enhancer-based biomarker of resistance to anti-PD-1 and suggest enhancer blockade in combination with ICB as a potential strategy to improve responses.

## INTRODUCTION

In recent years, there has been tremendous progress in melanoma immunotherapy, including the FDA approval of anti-CTLA-4 antibodies (in 2011) and anti-PD-1 antibodies (in 2014). Though response rates for monotherapy with these agents are modest (∼15% for anti-CTLA-4 and ∼44% for anti-PD-1), a subset of responses are often durable (Brahmer et al., 2012; Hodi et al., 2010; Schadendorf et al., 2013; Topalian et al., 2012), with 2-year survival rates up to 43% among patients who receive anti-PD-1 monotherapy and a 10-year survival rate of ∼20% for those who receive anti-CTLA-4 monotherapy (Topalian et al., 2012; Topalian et al., 2014). Response rates are also significantly increased by combination anti-PD-1/anti-CTLA-4 therapy (Postow et al., 2015). However, a significant proportion of patients still do not achieve clinical response, and exhibit severe toxicity (Postow et al., 2015). Therefore, there is a critical unmet need to identify biomarkers that predict response or resistance to immune checkpoint blockade (ICB)—either as monotherapy or in combination—and to identify actionable strategies that will enhance the effectiveness of these potent therapies in the patients most likely to benefit.

The epigenome consists of an array of chromatin modifications, including DNA methylation and histone marks, which collectively form a dynamic state that is referred to as a “chromatin state”. The nature of chromatin states and their impact on associated genomic loci are determined by their constituent histone or DNA modification marks (Lee and Young, 2013). For example, the presence of the H3K27me3 mark (trimethylation of lysine 27 on histone H3) in promoters is associated with transcriptional repression, whereas H3K4me3 (trimethylation of lysine 4) is associated with transcriptionally active promoters. H3K4me1-modified and H3K27Ac- modified nucleosomes are present only at enhancer elements, whereas the presence of H3K79me2 or H3K36me3 coincides with transcribed regions (Barski et al., 2007). Thus, profiles of histone modification marks generate a comprehensive map of the epigenome.

Recent data indicate that responsiveness to ICB therapy may be associated with specific epigenetic processes. For example, regulation of histone modifications by HDAC, EZH2, or KMT2D has been proposed to modulate either response to these agents or antitumor activity of immune cells (Maitituoheti et al., 2020; Peng et al., 2015; Wang et al., 2020; Woods et al., 2015). However, there is insufficient understanding of the epigenome content of ICB-sensitive and ICB-resistant cases. Furthermore, whether specific patterns of chromatin modification states are associated with response to ICB has not been systematically investigated. As chromatin modification states are stable and heritable, specific patterns of chromatin modification states can potentially be used as biomarkers for ICB response (Mulero-Navarro and Esteller, 2008).

By generating epigenome profiles of 36 melanoma samples treated with ICB at MD Anderson Cancer Center (MDACC), followed by validation in an independent cohort of 20 melanoma samples treated with ICB at Massachusetts General Hospital (MGH), we demonstrate that an enhancer signature of 437 genomic loci in pre-treatment samples can predict non-response of melanoma to ICB. Enhancer gains in non-responders were observed on a number of resistance-driving genes, and enhancer-blocking bromodomain inhibitors synergized with anti-PD-1 antibodies in pre-clinical models. Altogether, we identify enhancer gains as an important epigenetic mechanism driving resistance to anti-PD-1 therapy in melanoma, which could be leveraged for biomarker development or novel therapeutic combinations.

## RESULTS

To directly address whether epigenomic changes are associated with response to ICB therapy, we mapped chromatin state patterns in 36 metastatic melanoma samples from patients treated with nivolumab or pembrolizumab (anti-PD-1 antibodies) at MDACC (**Fig. 1A** and **Table S1**). Response in these patients was documented using RECIST criteria, which identified 4 samples from patients who achieved complete response, 4 with partial response, 5 with stable disease, and 23 with progressive disease in response to ICB therapy (**Fig. S1A-S1B**). Overall, 13 samples from patients with complete or partial response or stable disease were annotated as “responders (R)” and the 23 samples from patients with progressive disease were labeled as “non-responders (NR)” (**Fig. S1A-S1B**). Samples were collected at 3 timepoints: 1) pre-treatment (n = 17), 2) on-treatment (n = 4), and 3) post-treatment (n = 15).

**Figure 1:**
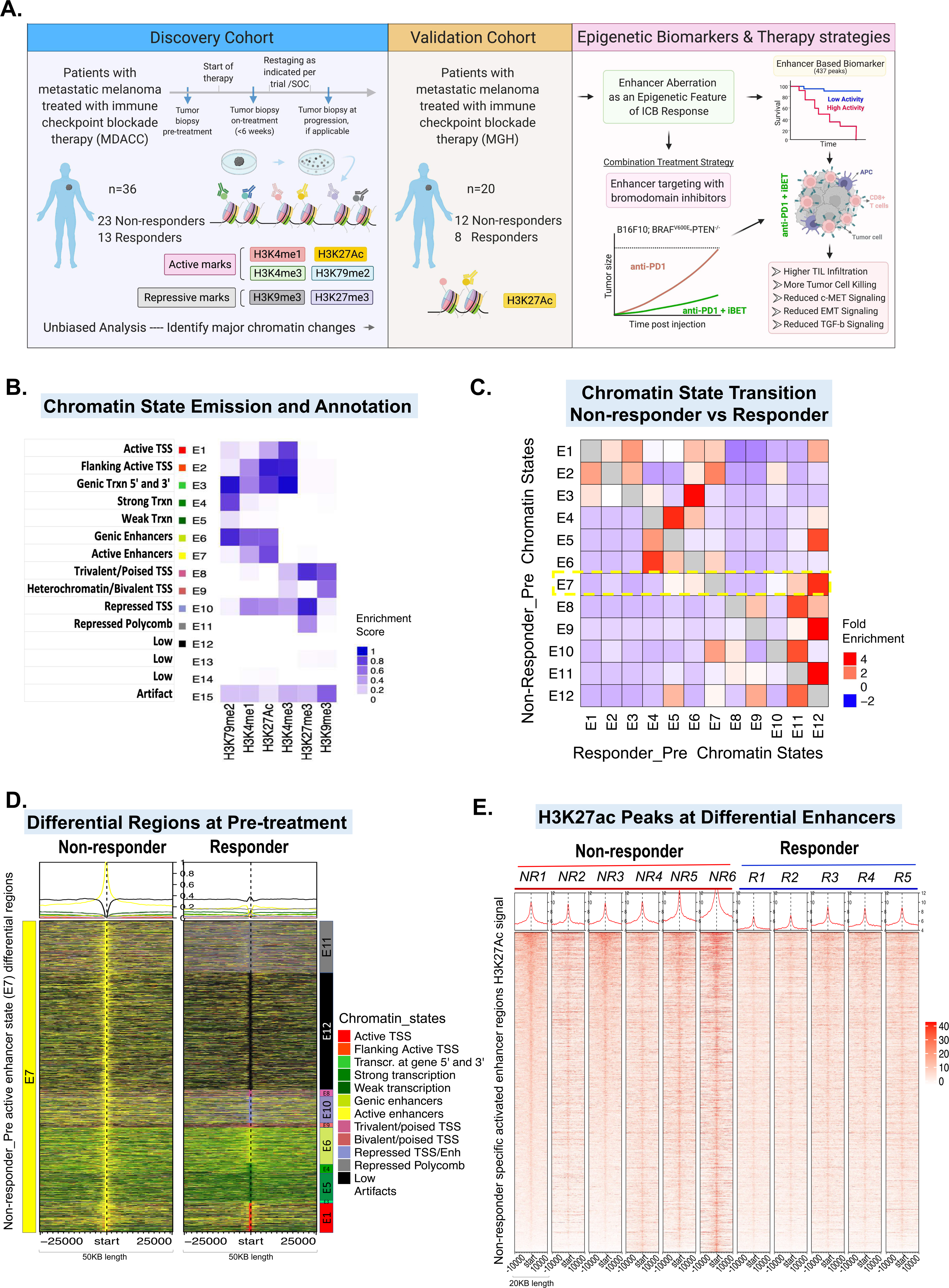
Comprehensive epigenome profiling of anti-PD-1–treated melanoma patients identify enhancer set as predictive biomarker of non-response to ICB. **A.** Schematic diagram describing the approach and main findings of the study. **B.** Emission parameters of the 15-state chromatin state model defined using ChromHMM on ChIP-Seq data for 6 histone modification marks (shown on x-axis) in the discovery cohort from MDACC (n = 36). Annotations on the left are derived from the relative enrichment of different histone marks and genomic distribution of the loci in that particular state (Fig. S1D). The intensity of the color in each cell reflects the frequency of occurrence of that mark in the corresponding chromatin state on the scale from 0 (white) to 1 (blue). **C.** Heatmap showing the fold enrichment of chromatin state transitions between responder and non-responder pre-treatment samples for the 15-state model defined by the ChromHMM. Color intensities represent the relative fold enrichment. Yellow box points to switches in active enhancer state E7 in non-responder to rest in responder. Diagonal is grayed to highlight non-self state transitions. **D.** Heatmap of chromatin state intensities for 31,155 loci that show switch from active enhancer state E7 (yellow) in non-responder pre-treatment samples (left) to any other state in responder pre-treatment samples (right) as shown by colors for each state. Note the high percentage of non-responder active enhancer state E7 transitioning to low or repressed states E12 (black), E11 (gray), or E10 (purple) in responders. p-value presented is for a 2-tailed Student *t*-test. **E.** Heatmap for H3K27ac mark in 24,862 peaks corresponding to 21,924 bins with active enhancer states that shows consistent depletion in responder pre-treatment samples (right 5 samples) compared with non-responder pre-treatment samples (left 6 samples). Enhancers are shown in a 20-kb window centered on the middle of the enhancer in non-responder and responder pre-treatment samples.

To identify chromatin state patterns, we profiled 6 reference histone modifications that mark promoter (H3K4me3), enhancer (H3K4me1 and H3K27Ac), transcribed (H3K79me2), and repressed (H3K27me3 and H3K9me3) states using high-throughput ChIP-sequencing methodology (Garber et al., 2012; Rai et al., 2015) in all 36 samples, generating 148 chromatin maps (**Fig. S1C**). This approach is similar to that utilized by ENCODE (Consortium et al., 2012) and NIH Roadmap projects (Bernstein et al., 2010) to determine basic epigenome maps in normal tissues and cell lines. As histone modifications exert their function in a combinatorial fashion, we identified such chromatin states using the ChromHMM algorithm (Ernst and Kellis, 2012). A model of 15 chromatin states was chosen for more in-depth interrogation into the biology of chromatin in anti-PD-1 response, as it presented sufficient resolution for biological interpretation (**Fig. 1B and Fig. S1D**). Annotation of these states based on the content of histone marks and their genomic locations revealed the presence of active promoter (E1, E2, E3), active enhancer (E6, E7), transcribed (E4, E5), polycomb-enriched (E11), heterochromatin/bivalent (E9), poised (E8, E10), and low (E12, E13, E14; merged as E12 afterwards) states (**Fig. 1B**).

### Chromatin state transitions between sensitive and resistant lesions

We first identified chromatin state differences between pre-treatment samples belonging to the responsive (R) and non-responsive (NR) groups. To this end, we consolidated chromatin states using epilogos (see **Methods**) and computed transitions in these states between the responder and non-responder samples (**Fig. 1C**). The most notable transition was from the active enhancer state E7 in non-responder samples to low (E12), polycomb (E11), or repressed (E10) states in responders, based on the number of switching bins in the responder and non-responder groups (**Fig. 1D**). We identified 31,555 bins (1-kb segments) that showed transitions between active enhancer state E7 in non-responsive samples to low, repressive states E10, E11, and E12 in responsive samples (**Fig. 1D**). These differences in active enhancer states showed significant, yet modest, changes in corresponding gene expression (**Fig. S1E**). Observed differences in active enhancer chromatin state signals on these loci were also recapitulated when only H3K27ac signals were examined. H3K27ac signal was decreased in 24,862 peaks corresponding to 21,924 bins with active enhancer states in pre-treatment samples from the responders compared to those from the non-responders (**Figs. 1E, S1F**). Average intensity of H3K27ac on these enhancers also showed a drastic increase in the non-responders compared to the responders, whereas the average intensity for H3K27me3 occupancy on these enhancers was significantly increased (**Fig. S1G**). We also noted that loci harboring active enhancer (E7) state in pre-treatment R samples, but not in pre-treatment NR samples were enriched surrounding genes involved in T cell function suggesting higher lymphocyte infiltration in responder samples (**Figs. S1H**).

### An enhancer signature predicts response to anti-PD-1 therapy in melanoma

To establish multiple independent lines of evidence supporting a concrete set of epigenomic correlates of ICB resistance, we collected an independent cohort of 22 melanoma samples from the MGH biobank and generated H3K27ac ChIP-seq data (**Fig. 1A and Table S1**). To make our MDACC and the MGH datasets jointly analyzable, we defined a common metric that could be used across both cohorts by using MAnorm (Shao et al., 2012) to calculate a log2 ratio of read densities (M-value) between ChIP and a whole-cell extract control that was adjusted for the average log2 read density at all peaks (**Figs. S2A-S2B**). This allows any 2 peak regions to be compared on the same scale between the two distinct cohorts by accounting for variable total read depth at peak regions of interest. Using IDR (Irreproducible Discovery Rate) analysis (see **Methods**), we identified a subset of 84,317 out of 244,472 peaks as reproducible between the MDACC and MGH cohorts (**Fig. 2A**). These peaks were enriched in various functional classes, including promoter, intron, and intergenic areas (**Fig. S2C**).

**Figure 2:**
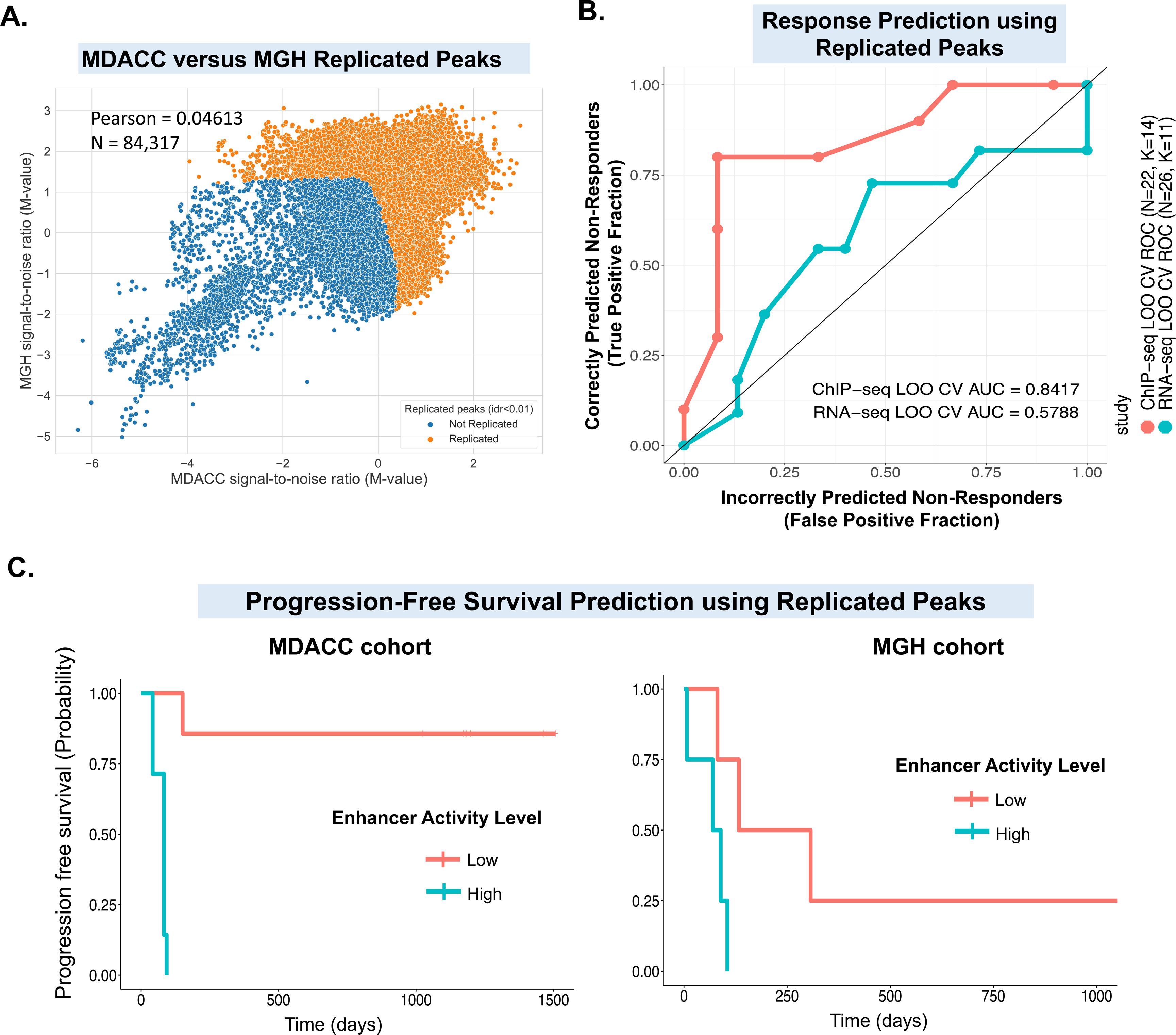
Validation of enhancer signature’s prediction of nonresponse in an independent cohort. A. Average M-values for MGH vs. MDACC cohorts with IDR (Irreproducible Discovery Rate) status <0.01. Individual points represent averaged M-value across the MDACC cohort (x-axis) and across the MGH cohort (y-axis). Color denotes whether a particular peak was flagged by IDR. B. Receiver operating characteristic (ROC) of random forest trained predictive models using leave-one-out cross-validation across N=22 pre-treatment ChIP-seq samples and N=26 pre- treatment RNA-seq samples. The features within each cross-validation fold were determined by finding the set of replicated peaks or genes across the MDACC and MGH cohorts and by computing the set of intersecting peaks or genes that were nominally significant in the training set within both cohorts. We analyzed a total of 23,457 RNA-seq genes and 84,317 ChIP-seq peaks. This was used to train a random forest within each training set with K=5 to K=20 trees. ROC and auROCs were derived from the best performing random forest classifier. The ROC curve and auROC was formed by concatenating predictions from the N=22 ChIP-seq and N=26 RNA-seq cross validation folds. C. Kaplan-Meier plots for progression-free survival of patients in MDACC (left) or MGH (right) cohorts for 29 out of the 32 peaks which offered worse prognosis as a result of increased peak signal. The normalized ChIP activity values were studentized across the MDACC cohort, then the median value was used to determine the high-activity vs. low-activity groups. There was a total of 8 patients (4 low enhancer activity, 4 high enhancer activity) in the MGH group and a total of 14 patients (7 low enhancer activity, 7 high enhancer activity) in the MDACC group. The peaks were selected using a p=0.05 cutoff for the Cox proportional hazards test.

Next, we subjected the M-values of this subset of peaks to differential peak calling via *limma* (Ritchie et al., 2015) in each cohort independently. We identified 5174 MGH and 8291 MDACC pre-treatment peaks whose activity was significantly (p < 0.05) different between responders and non-responders (**Table S2**). To identify a replicated peak set, we intersected differentially enriched peaks within both the MDACC and MGH cohorts to determine whether this set exhibited statistically significant enrichment, above the null expectation. Only the pre-treatment comparisons exhibited a significant enrichment in the number of replicated peaks (p = <2.2e-16, one-sided exact binomial test), and 437 peaks were doubly significant in both the MDACC and MGH pre-treatment comparisons. We also noted excess enrichment in the signal from the MDACC cohorts in both the pre-treatment (**Fig. S2D**) and on-treatment (**Fig. S2E**) comparisons. We also generated RNA-seq data on 44 ICB-treated melanoma samples consisting of 26 pre-treatment (14 NR and 12 R), 10 on-treatment (6 NR and 4 R) and 8 post-treatment (6 NR and 2 R) samples from both MDACC and MGH cohorts. Here we noted 588 differentially expressed genes (DEGs) between NR and R at pre-treatment stage (**Table S2**).

Next, to identify a subset of enhancers with predictive ability for patient response, we concentrated on the pre-treatment significant H3K27ac peak set overlapping between the MDACC and MGH cohorts. We utilized the 437 replicated peaks as a feature set in a cross-validation setting and trained 2 random forest models: one in which the MDACC cohort was designated as the training set and the MGH cohort the testing set, and vice versa. The results were combined into a single receiver operator characteristic (ROC) for evaluation. We also evaluated the area under the ROC curve (AUC) as a measure of model performance. Using the 437 peaks, we were able to achieve an AUC of 0.9 (**Fig. S2F**). However, in this analysis, features were determined by the union of the two datasets making it prone to potential data leakage between the training and testing cohorts. To prevent this issue, we performed leave-one-out (LOO) cross validation (CV) on N=22 pre-treatment ChIP-seq samples and N=26 pre-treatment RNA-seq across the two cohorts. To generate the features, within each cross-validation fold (N=21 ChIP-seq training samples, N=25 RNA-seq training samples), we repeated our replicated peak calling procedure by finding the overlapping peaks with nominal p<0.05 in both the MDACC and MGH cohorts for each CV fold. We tested a total of 23,4457 RNA-seq genes and 84,317 ChIP-seq peaks. These features were then used to train a random forest classifier with K=5 to K=20 trees on the training set, and subsequently tested on the N=1 testing set to construct the ROC across the 22 ChIP-seq and 26 RNA-seq CV folds. We took the highest performing classifiers for the ChIP-seq and RNA-seq separately and reported their performance. We observed that epigenomic features are moderately predictive of immunotherapy response, with an AUC of 0.842 (**Fig. 2B**). On the other hand, RNA-seq features showed much less predictive ability (AUC = 0.579) when evaluated using the same feature discovery framework described above (**Fig. 2B**). This relationship holds when the aforementioned approach is applied only on the pre-treatment samples for which both RNA-seq and ChIP-seq data are available (**Fig. S2G**). Our enhancer based classifier also performed better than prior biomarkers based on RNA expression patterns, tumor mutation burden (TMB), or histopathological features (Auslander et al., 2018; Johannet et al., 2021; Shi et al., 2020; Yan et al., 2020) (**Fig. S2H**). We further examined TMB in MDACC cohort by generating and analyzing WGS data from 34 samples and in MGH cohort by analyzing WES data from 8 samples, but failed to observe significant difference in mutation burden between R and NR patients (**Fig. S2I**). We also utilized the TMB data from pre-treatment samples as a predictive feature for response in the MDACC cohort. In LOO-CV across N=13 MDACC samples with both H3K27ac and TMB data, we observed incorporating TMB data along with differential H3K27ac peaks (AUC=0.7143) as features to a random forest classifier with K=20 trees resulted in a slightly increased AUC compared to only using differential H3K27ac peaks alone (AUC=0.6905) **(Fig. S2J).**

We next assayed to what extent these 437 peaks stratified progression-free survival in our clinical cohort. To do so, we performed Cox proportional hazards regression with M-values as the design matrix which showed that 32 out of the 437 peaks significantly stratified survival in both the MGH and MDACC cohorts. As a result of increased peak signal, 29 out of the 32 peaks offered worse prognosis (**Fig. 2C)**, whereas 3 out of 32 peaks offered better prognosis **(Fig. S2K**). Our results show that a distinct set of epigenomic peaks are significantly associated with treatment response and survival stratification in 2 independent cohorts, making these peaks optimal targets for follow-up prognostic studies.

### Enhancer activation targets genes contributing to anti-PD-1 resistance

Do these differential enhancers between non-responders and responders play functional role during evolution of ICB resistance? To address this question, we first sought to identify the gene targets of NR- or R-specific enhancers by overlapping them with the enhancer-promoter (E-P) annotation. As enhancers activate their target gene expression by looping onto the promoter, the E-P annotation was predicted using in-house H3K27ac HiChIP data from one of the short-term melanoma culture (STC2765 which is derived from anti-PD-1 non-responder melanoma tumor) and from a prior study using 935 samples, covering a major fraction of human cell and tissue types (ENCODE + Roadmap or FANTOM5)(Cao et al., 2017) (see **Methods**). This identified 1318 gene targets of 966 reproducibly enriched enhancers (false discovery rate [FDR] < 0.1) in non-responsive samples (**Table S2**). To dissect whether enhancer peaks were derived from melanoma cells or tumor-infiltrating T cells (TILs), we overlapped the replicated enhancer peaks (562 NR-specific and 161 R-specific) with in-house H3K27ac ChIP-seq data on short-term melanoma cultures (STCs) from 10 patients (Terranova et al., 2021) and cognate TILs derived from 8 of them (**Fig. 3A and Table S3**). Pathway analysis of target genes of melanoma tumor cell enriched NR-specific enhancers showed MAPK pathway, Epithelial-to-Mesenchymal transitions (EMT), TGFβ pathway among others (**Fig. 3A),** some of which have been previously implicated in immune evasion and immunotherapy resistance (Mariathasan et al., 2018; Terry et al., 2017).

**Figure 3:**
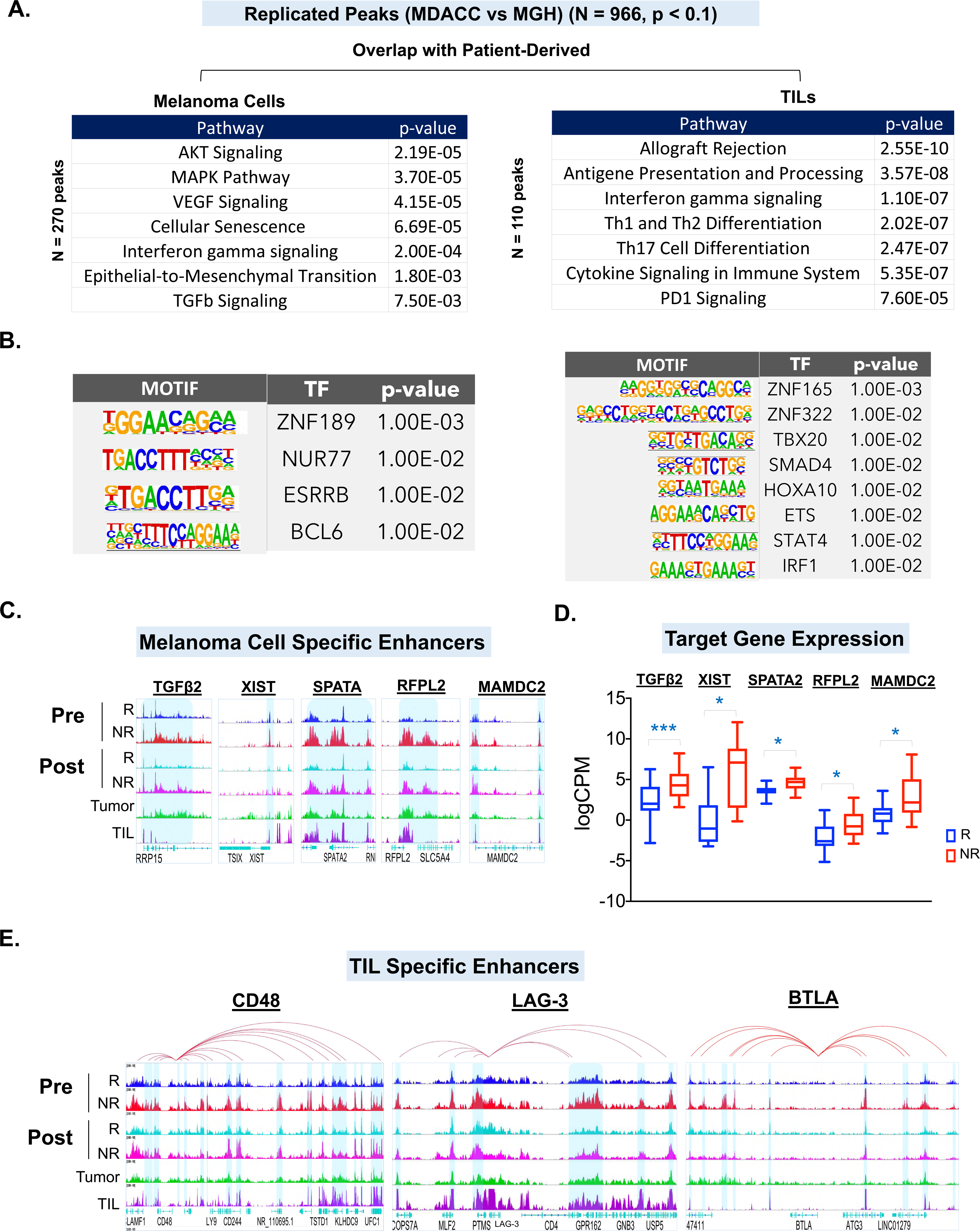
Enhancer activation marks key immune resistance-associated genes in anti-PD- 1 non-responders. A. List of significantly enriched pathways in genes targeted by replicated H3K27ac peaks (n=966, p < 0.1) that overlap with those from isolated melanoma cells (n=270, left) or TILs (n=110, right) enhancer peaks. B. List of significantly enriched transcription factor (TF) motifs in replicated H3K27ac peaks that overlap with those derived from isolated melanoma cells (n=270, left) or TILs (n=110, right). C. IGV (Integrated Genomic Viewer) snapshot of aggregate H3K27ac profiles around TGFβ2, XIST, SPATA2, RFPL2 and MAMDC2 genes in MDACC cohort non-responder (NR) samples, responder (R) samples, isolated melanoma STCs, or isolated TILs. Highlighted regions show enrichment of H3K27Ac enhancer peaks in NR samples compared to R samples. D. Box plot showing mRNA expression level of TGFβ2, XIST, SPATA2, RFPL2 and MAMDC2 genes in NR and R pre-treatment samples from both cohorts. In the box plot, the bottom and top of the rectangles indicate the first quartile (Q1) and third quartile (Q3), respectively. The horizontal lines in the middle signify the median (Q2), and the vertical lines that extend from the top and the bottom of the plot indicate the maximum and minimum values, respectively. E. IGV snapshot of aggregate H3K27ac profiles around CD48, LAG-3, and BTLA genes in MDACC cohort NR samples, R samples, isolated melanoma STCs, or isolated TILs. The red line loops depict E-P interactions identified from H3K27ac HiChIP data from STC2765 cells and/or previously predicted E-P networks (Cao et al., 2017). Highlighted regions show enrichment of H3K27Ac enhancer peaks in NR samples compared to R samples.

These genes included known regulators of anti-tumor immune response such as NOTCH1, AKT1, TGFβ2, USP22, MYC, MITF, c-MET (**Fig. S3A**)(Batlle and Massague, 2019; Casey et al., 2016; Li et al., 2020; Meurette and Mehlen, 2018; Papaccio et al., 2018; Rogel et al., 2017; Wiedemann et al., 2019). Motif enrichment analysis (HOMER) provided insight into TFs that are known (such as NUR77, STAT4, IRF1) or unknown (such as ZNF189, ESSRB, BCL6, TBX20, SMAD4) to contribute to immune evasion or ICB resistance within the melanoma or TILs (**Fig. 3B**). Recent study of whole-exome and transcriptome meta-analysis of over 1,000 patients treated with ICB revealed that CXCL9/CXCL13 are the strongest predictors of response (Litchfield et al., 2021), consistently we also noted enhancer enrichment nearby these two genes in responder tumors (**Fig. S3B**). In NR samples, we also detected enhancer gains on TGFβ, PI3K-AKT and angiogenesis pathway genes that are known to cause systemic immunosuppression (**Figs. S3C-S3E**)(Fukumura et al., 2018). Finally, we identified other potentially novel regulators of anti-tumor immune response such as FAM20C, RFPL2, MAMDC2, SPATA2 in melanoma cells (**Figs. 3C, S3F-S3G and Table S3**) (Lee et al., 2020; Schlicher et al., 2016; Xu et al., 2021). Integration of enhancer gains with gene expression data showed concomitant upregulation of gene expression of a subset of enhancer-target genes at the pre-treatment stage (**Figs. 3D, S3I**).

Target genes of TILs-enriched enhancers (from 437 replicated H3K27ac peaks) were enriched in allogenic transplant, interferon signaling and other pathways which are known to play important roles in T cell differentiation and anti-tumor activity (**Fig. 3A)**. Overlap of enhancers enriched in NR pre- or post-treatment tumors with those in isolated TILs identified genes in multiple categories: 1) known inhibitors of T cell activity such as CISH (Palmer et al., 2015); 2) important inhibitory checkpoint receptors, such as LAG-3 (Joller and Kuchroo, 2017) and BTLA (Watanabe et al., 2003), or their key partners, such as CD48 and CEACAM-1 (required for function of TIM-3 (Huang et al., 2015)); 3) genes known to mediate key interactions with antigen-presenting cells or tumor cells CD244 and HVEM (Wherry and Kurachi, 2015); 4) transcription factors mediating T-cell exhaustion such as NR4A1(Chen et al., 2019)(**Figs. 3E, S4A-S4C**); 5) potential novel regulators of T cell function such as FKBP3, LGALSL, LARP1, CEBPβ and KLF6 (**Fig. S4D**). Overall, these data suggest that replicated enhancers enriched in pre-treatment NR samples activate multiple resistance mechanisms in the melanoma cells as well as infiltrating T cells.

### cMET Enhancers play functional role in mediating anti-tumor killing

To gain a deeper insight into functional role of enhancer gains in ICB response biology, we focused on c-MET which showed increased enhancer peaks and associated gene expression in NR at pre- or post-treatment stage (**Figs. 4A-4B, S4E-S4F**). The c-MET locus harbored multiple distal enhancers that were present in NR tumors, but not in R tumors, and the HiChIP data provided evidence for looping between 4 distal enhancers (E1, E2, E3, and E4) and gene body/transcription start site (TSS) (**Fig. 4A, 4C**). These enhancers were also present in STC2765 melanoma cells as suggested from overlapping H3K27ac peaks (**Fig. 4A**). Consistently, c-MET expression was localized to melanoma cells when published single cell RNA-Seq data was queried (**Fig. S4G**) (Tirosh et al., 2016). Silencing of these enhancers using specific gRNAs and dCas9-KRAB (Klann et al., 2017) significantly reduced expression of the c-MET gene in STC2765 cells **(Fig. 4C**). The cell lines with dCas9-KRAB–mediated enhancer suppression also showed increased tumor killing by autologous T cells (TIL2765 that were derived from the same tumor as STC2765) in a co-culture assay, thus demonstrating enhancer functionality (**Fig. 4D**). Consistently, treatment with a c-MET inhibitor (Crizotinib) also showed enhanced T cell–mediated killing of STC2765 cells by TIL2765 (**Fig. 4E**). These data provide c-MET enhancers as an example of functional enhancer elements that contributes to immune evasion process during anti-PD-1 treatment. Taken together with enhancer activation surrounding numerous regulators of anti-tumor immune response (Fig. 3), these data suggest that activation of enhancers could be a key epigenetic mechanism for activation of many regulators and cellular processes that promote resistance to ICB therapy.

**Figure 4:**
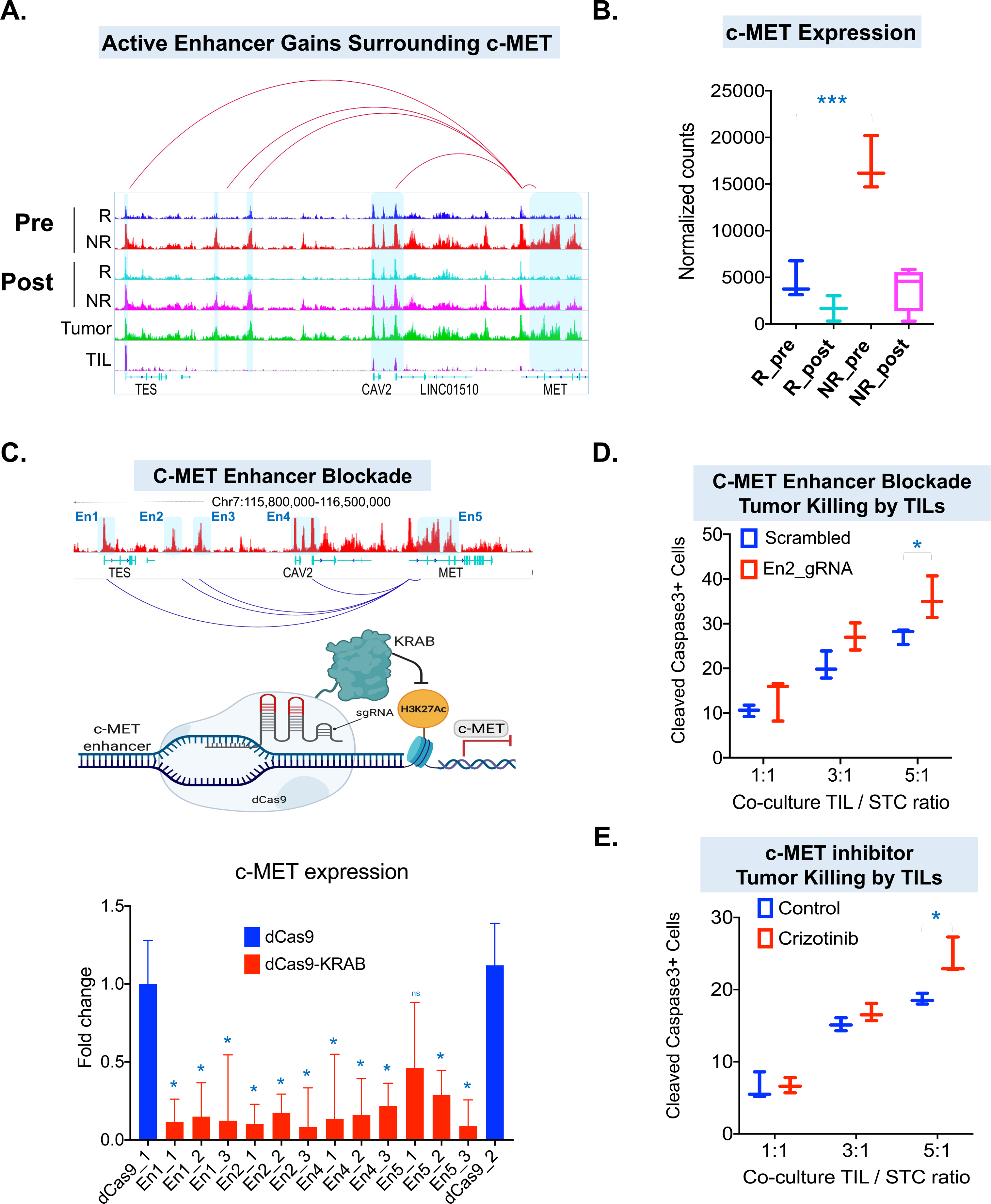
Enhancer activation of c-MET contributes to non-response to ICB. **A.** IGV snapshot of aggregate H3K27ac profiles around c-MET gene in NR samples, R samples, isolated melanoma STCs, or isolated TILs. The red line loops depict E-P interactions identified from H3K27ac HiChIP data from STC2765 cells and/or previously predicted E-P networks (Cao et al., 2017). Highlighted regions show enrichment of H3K27Ac enhancer peaks in NR samples compared to R samples. **B.** Box plots showing normalized RNA counts for c-MET gene between NR and R samples at pre- and post-treatment stages. **C.** Top, genomic locations of c-MET enhancers (En1 through En5) and HiChIP-derived E-P loops. Middle, schematic of dCas9-KRAB mediated repression of c-MET enhancer. Bottom, bar plot showing fold change of gene expression for c-MET gene upon targeting of enhancers by specific gRNAs as indicated. **D-E.** Percentage of cleaved caspase-3 positive STC2765 post co-culture with autologous TIL2765 at 3 different ratios of TIL:STC ratio (1:1, 3:1 and 5:1). In panel **D**, STC2765 cells harboring dCas9- KRAB and control or c-MET enhancer gRNAs were used as target cells, whereas in panel **E**, parental STC2765 cells were treated with c-MET inhibitor crizotinib (for 24 hrs at 2μM) before and during co-culture. In (**B**) and (**D-E**) box plots, the bottom and top of the rectangles indicate the first quartile (Q1) and third quartile (Q3), respectively. The horizontal lines in the middle signify the median (Q2), and the vertical lines that extend from the top and the bottom of the plot indicate the maximum and minimum values, respectively.

### Enhancer reprogramming during ICB treatment

We next sought to define dynamics of chromatin states as patients progress or respond to anti-PD-1 therapy by computing chromatin state transitions between pre- and post-treatment samples. In responders, we primarily observed transitions of active states in pre-treatment to repressed states in the post-treatment. On the other hand, transitions in the non-responder samples were distributed more evenly between repressive and active states (**Fig. S5A**). To determine the reprogramming of active enhancers during the treatment stage, we computed the chromatin state transition of active enhancer state E7 between post-treatment and pre-treatment samples (**Fig. S5B**). Seven clusters were identified based on the transition of enhancer states, of which Cluster 1 enhancers gained repressive states or lost the active enhancer marking, whereas Cluster 4 enhancers remained in active enhancer state even at the post-treatment stage (**Fig. S5B**). Cluster 1 enhancers were enriched in VEGFA, autophagy, and HIF1 signaling, including VEGFA, RUNX3, and AKT2 genes (**Figs. S5C-S5D and Table S4**). Unaffected Cluster 4 enhancers were enriched in TGFβ, PI3K/AKT/mTOR signaling pathways, AHR, and oxidative stress pathways, including genes such as the TGFβ and LOXL4 (**Figs. S5E-S5F**). These observations provide better understanding of the dynamics of enhancer states on specific pathways during anti-PD1 treatment.

### Combination of bromodomain inhibitors with anti-PD-1 enhances response in mouse melanoma models

Since enhancer activation marks multiple genes that regulate resistance to anti-PD-1 antibodies, we reasoned that inhibitors of acetylation-reader bromodomain, which relay the signal from the enhancers, could be used as an umbrella approach to target many resistance mechanisms at once along with anti-PD-1 therapy to enhance its efficacy. BRD4 (bromodomain containing protein 4) has been previously implicated as a major reader of H3K27ac on active enhancers that acts with other transcriptional regulators to activate or enhance gene expression(Kanno et al., 2014). We noted higher BRD4 levels in metastatic melanoma in comparison to primary tumors in The Cancer Genome Atlas (TCGA) skin cutaneous melanoma (SKCM) dataset (**Fig. 5A**). Importantly, the tumors harboring higher levels of BRD4 survived poorly in comparison to those harboring lower levels of this protein (**Fig. 5B**). Similar trend for BRD4 (and other family members) expression with progression-free survival was also observed in Schadendorf cohort (Liu et al., 2019) of advanced melanoma patients treated with anti-PD-1 (but without prior anti-CTLA-4 treatment) (**Figs. 5C, S6A**). Similar to previous reports in ovarian and triple negative breast cancers (Jing et al., 2020; Zhu et al., 2016), we also observed positive correlation between BRD4 expression and PDL1 expression (**Fig. 5D**). These clinical validations of the BRD4 manifest it as an optimal therapeutic target in melanoma. Treatment of tumors generated by transplantation of murine melanoma cell lines BP [from the Bosenberg model (Dankort et al., 2009)] and B16-F10 with the combination of iBET-762 and anti-PD-1 antibody significantly reduced tumor growth at doses that failed to generate much response when used as monotherapy (**Figs. 5E-5F**). Profiling of CD8+ T cells in these experiments revealed increased infiltration of these cells upon combination treatment in comparison to monotherapy (**Fig. 5G, S6B**). Consistently, we noted a modest negative correlation between BRD4 expression and infiltrating tumor cells in the TCGA cohort (**Figs. 5H, S6C)**. In addition, treatment of STC2765 cells with bromodomain inhibitors increased the TIL2765-mediated killing in a co-culture assay (**Fig. 5I)** and increased the MHC class I expression on tumor cells (**Fig. S6D)**.

**Figure 5:**
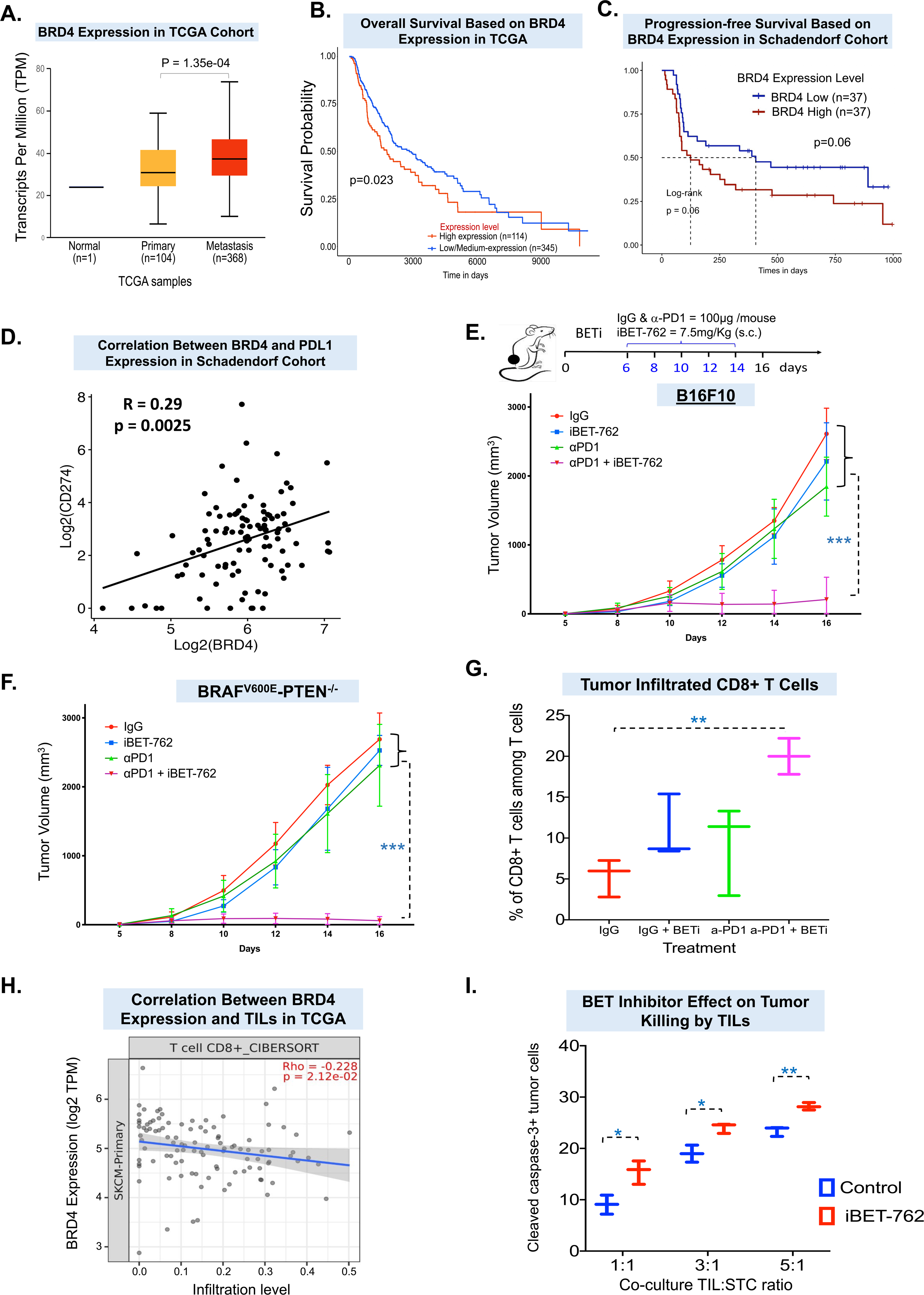
Targeting enhancers using bromodomain inhibitors in combination with anti-PD- 1 antibody confers synergistic tumor growth reduction. **A.** Box plot showing mRNA expression level of BRD4 in TCGA normal, primary, and metastatic melanoma patient samples. Metastatic melanoma patient samples display significantly higher expression in comparison to primary tumors. **B.** Kaplan-Meier plot of survival of melanoma samples in the TCGA, comparing overall survival between groups with high BRD4 expression (n=114) and low BRD4 expression (n=345). The log-rank (Mantel-Cox) *p* value was used to assess the significance of difference in survival. **C.** Kaplan-Meier plot of progression-free survival of anti-PD-1 treated (without prior anti-CTLA4 treatment, Schadendorf cohort) patients with high versus low BRD4 expression (split by median). The log-rank (Mantel-Cox) *p* value was used to assess the significance of difference in survival. **D.** Scatter plot showing the positive correlation between BRD4 and PD-L1 expression in Schadendorf cohort (Spearman’s rank test). **E.** Top: Schematic for mouse treatments. Bottom: Tumor growth curves for mice in 4 treatment categories: 1) IgG alone (100 μg/mouse), 2) anti-PD-1 antibody (100 μg/mouse), 3) bromodomain inhibitor iBET-762 (7.5 mg/kg) with IgG, or 4) iBET-762 with anti-PD-1 in B16-F10 cells. **F.** Tumor growth curves for BP cells derived from Bosenberg’s model (Tyr-Cre^ERT2^, BRAF^V600E^, PTEN^L/L^) upon treatment with the 4 different strategies shown in panel **E**. **G.** Graph showing the flow cytometry analysis results of infiltrated CD8+ T-cell percentages in tumors derived from experiment shown in panel **E**. **H.** Scatter plot showing the negative correlation between BRD4 expression (Log2TPM where TPM represents Transcripts per Million) and TIL infiltration score in TCGA melanoma cohort. **I.** Percentage of cleaved caspase-3 positive STC2765 cells post co-culture with autologous TIL2765 at 3 different ratios effector:target ratio (1:1, 3:1 and 5:1) when melanoma cells were treated with mock or iBET-762 (1μM) for 72 hrs. In (**A), (G)** and (**I**) box plots, the bottom and the top of the rectangles indicate the first quartile (Q1) and third quartile (Q3), respectively. The horizontal lines in the middle signify the median (Q2), and the vertical lines that extend from the top and the bottom of the plot indicate the maximum and minimum values, respectively.

### Bromodomain inhibitor combination with anti-PD-1 downregulates ICB-resistance pathways

To investigate the molecular mechanism underlying efficacy of bromodomain inhibitors and anti-PD-1 combination, we generated RNA sequencing-based transcriptome profiles and ChIP-Seq based genome-wide occupancy profiles for BRD4 and H3K27ac in the tumors from different groups of treatment in mice. Analysis of RNA-Seq data showed that genes overexpressed in the tumors treated with combination (iBET-762 plus anti-PD-1) versus control were associated with immune response, while repressed genes were associated with TGFβ, MYC and epithelial–mesenchymal transition (EMT) pathways (**Fig. 6A)**. Comparative analysis of H3K27ac ChIP-Seq data showed a significant decrease of average intensity of H3K27ac-marked enhancers in combination therapy versus vehicle control group while monotherapy showed intermediate effect (**Fig. S6E).** Integration of differentially enriched enhancers (DEEs) with DEGs showed loss of expression of a large number of genes (N = 714) in the combination treatment group in comparison to the monotherapy group (**Table S5**) or control (IgG) treated samples. While comparing these data with those from human patients (**Fig. 3**), we also noted reduced BRD4 and H3K27ac binding on enhancers for c-MET, TGFβ and genes belonging to PI3K-AKT-MTOR pathway, angiogenesis pathway (Fukumura et al., 2018) as well as immune checkpoint receptors in the combination treatment versus control groups. These data suggest that enhancer depletion may contribute to the decrease in tumor growth associated with combination treatment (**Figs. 6B, S6F-S6J**). Importantly, enhancer loss on many of these genes were associated with decreased gene expression in the combination treatment group (**Figs. 6C, S6G-S6I, S6K).** Hence, we extended the integration between DEGs and DEEs in mouse experiment to the gene targets of NR-enriched enhancers from patient samples. This revealed 107 genes with co-incident loss of expression, loss of binding of BRD4 and loss of H3K27ac active enhancer marks in combination iBET-762 plus anti-PD-1 treatment in comparison to the control treated group (**Fig. 6D**). These genes were enriched in WNT, TGFβ, epithelial-to-mesenchymal transition, and UV response pathways (**Figs. 6E, S6L**). Overall, these data provide evidence for enhancer-mediated activation of key resistance-driving genes/pathways as an epigenetic mechanism for resistance to ICB and demonstrate the need for clinical studies focused on the combination of enhancer-blocking agents and ICB to improve the response rate in melanoma and potentially other malignancies.

**Figure 6:**
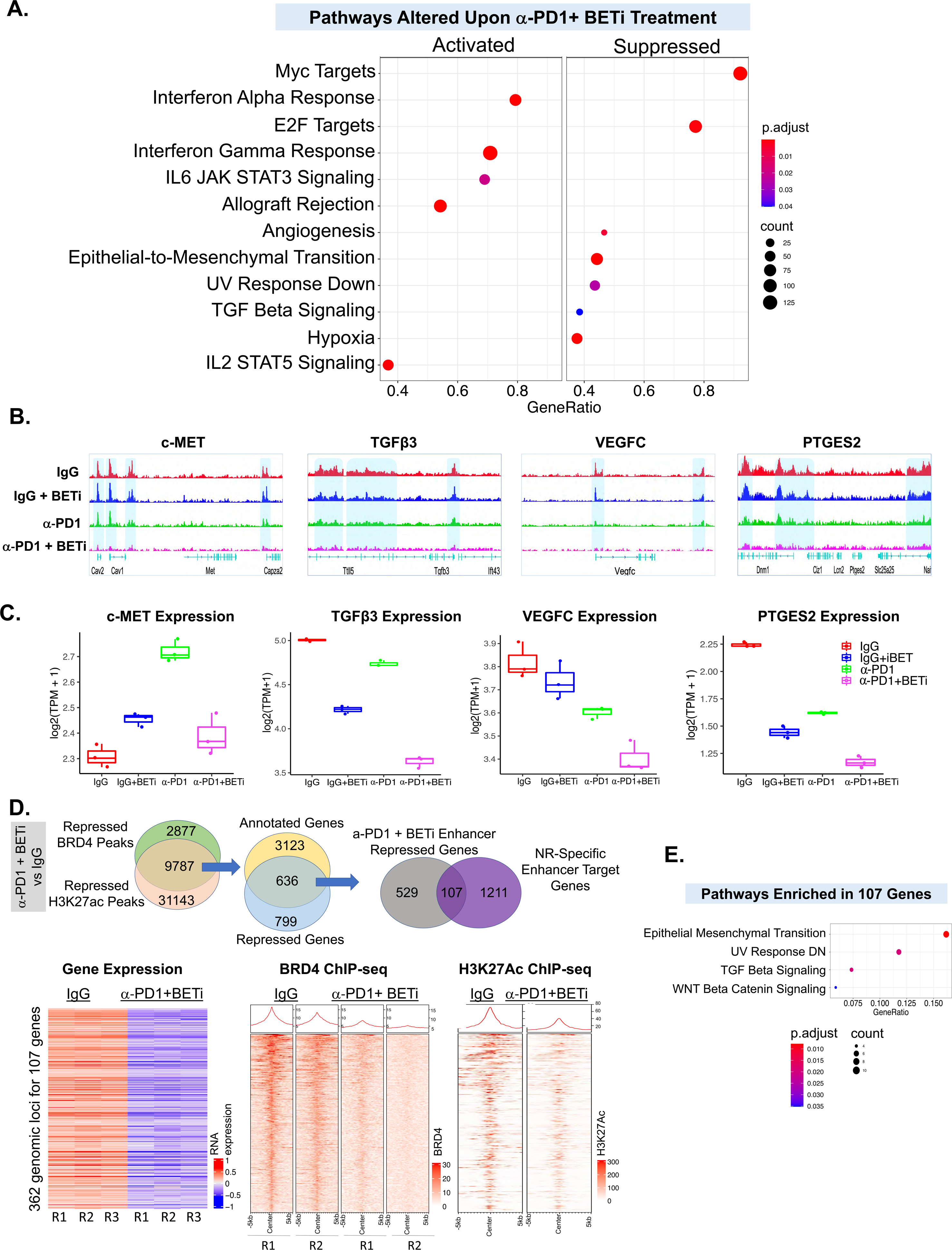
Molecular mechanism behind BRDi plus anti-PD-1 response. A. Dot plot showing significantly activated (left) and suppressed (right) pathways for differentially expressed genes in iBET-762 + anti-PD-1 treated tumors in comparison to anti-PD-1 treated ones. Dot size represents gene ratio, and colors represents adjusted p-values. B. IGV snapshot of aggregate BRD4 profiles around genes in c-MET, TGFβ3, VEGFC and PTGES2 in two different tumors belonging to the 4 treatment groups shown in Fig. 5E. Highlighted regions show loss of BRD4 peaks in iBET-762 + anti-PD-1 combination treatment samples compared to other treatment groups. C. Box plot for mRNA expression level (Log2TPM + 1) of genes shown in panel **B**. Each dot represents single sample. Colors represent 4 treatment groups shown in the plot. Bottom and top of the rectangles indicate the first quartile (Q1) and third quartile (Q3), respectively. The horizontal lines in the middle signify the median (Q2), and the vertical lines that extend from the top and the bottom of the plot indicate the maximum and minimum values, respectively. D. Top: Venn diagram showing the schematic of our approach to integrate mouse and human data. Bottom: Heatmaps for BRD4 and H3K27ac ChIP-Seq signal around differentially expressed genes overlapping between those in iBET-762 + anti-PD-1 versus IgG treated tumors and replicated non-responder–specific enhancers annotated genes. E. Pathway analysis (Hallmark) of 107 genes from human and mouse data overlap shown in panel D. Dot size represents gene ratio, and colors represents adjusted p-values.

## DISCUSSION

Our data help address two major clinical needs regarding ICB therapy in metastatic melanoma: 1) biomarkers that predict ICB response and 2) combination therapy strategies to improve the response to ICB. We observed that gains in enhancer activity on a set of genomic loci are associated with response to ICB and thus could potentially act as a predictive biomarker of response to ICB in metastatic melanoma. Our data also suggest causative roles for enhancer gains in non-response to ICB and supports the use of enhancer-blocking clinical agents in combination with anti-PD-1 as a potential strategy that can be tested in future clinical trials.

We identify an enhancer-based signature of 437 enhancers that could potentially be used as an epigenomic biomarker for non-response to ICB therapy in melanoma. Of these, 32 enhancers further predicted progression-free survival, suggesting the potential use of this epigenomic signature as a prognostic indicator. These signatures have the potential to be utilized alone or in combination with other genomic, transcriptomic, or immune features to generate a multi-omic signature to predict response or survival in patients on ICB therapy. Indeed, incorporation of other features such as tumor mutation burden (Goodman et al., 2017) may enhance this predictive potential of enhancer signatures. Other features, such as specific genetic features, such as PTEN deletion (Peng et al., 2016), IFNγR deletions (Gao et al., 2016), PBRM1 mutations, and KMT2D mutations (Wang et al., 2020), have been associated with response to ICB and could also be considered in combination with an enhancer-based signature to develop better prognostic biomarkers.

Our work suggests that pre-existing enhancer states could contribute to innate resistance to immune checkpoint therapy. Importantly, the overlap between the MDACC and MGH cohorts was highly significant at the pre-treatment stage, but not at on-treatment or post-treatment stage, suggesting that baseline chromatin states of the tumor are likely important drivers of ICB response and impact of ICB therapy on enhancer patterns of melanoma tumors significantly varies between different patients. Indeed, pre-existing chromatin states differ between individuals and, depending on their nature, could act as barrier to or facilitate activation of the tumor pathways that mediate immune recognition, immunogenicity or T cell–mediated killing (e.g., checkpoint receptors/ligands such as PD-L1, IFN/GAS/STING pathways, MHC expression or EMT) (Blank et al., 2005; Terry et al., 2017) (Bhat et al., 2017; Garcia-Lora et al., 2003; Kwon and Bakhoum, 2020). As an extension, adaptive resistance could also result from gain of enhancers on ICB resistant pathways during the course of treatment.

Bromodomain inhibitors serve as a useful tool for assaying whether enhancer blockade contributes to a specific phenotype. Here, combinations of bromodomain inhibitors enhance the activity of anti-PD1 suggesting that enhancer activation is likely a barrier to T-cell mediated killing of tumor cells. Indeed, integrative analysis of transcriptomic and epigenomic data between murine tumors treated with BRDi/anti-PD-1 combo and human anti-PD-1 tumors shows 362 enhancers (and their 107 target genes) as likely significant contributors to anti-PD-1 response. These enhancers target key pathways of WNT/β-catenin, EMT, TGFβ and UV response consistent with prior roles of some of these pathways in immune modulation (Luke et al., 2019; Mariathasan et al., 2018; Terry et al., 2017; Trujillo et al., 2019). These enhancers/genes could potentially serve as pharmacodynamic markers for BRDi in future clinical/pre-clinical studies.

Overall, our studies support further prospective clinical investigation into the utility of these enhancer signatures in predicting response to immunotherapy, offering prognostic information, and informing combinatorial clinical trials facilitated by cutting-edge epigenomic tools. Finally, our data suggest a need for future clinical studies to test potency of the BRDi or other enhancer blocking inhibitors [such as CDK9/7(Kwiatkowski et al., 2014; Morales and Giordano, 2016)] in combination with immune checkpoint inhibitors.

## METHODS

### Patient samples

Tissue samples from metastatic melanoma patients were collected and viably frozen as part of an IRB-approved tissue banking protocol at MDACC and MGH. All patients signed written informed consent prior to having the sample collected. All patients received either pembrolizumab or nivolumab as the anti-PD-1 therapy for their metastatic melanoma. Thirty-six melanoma tumor samples (17 samples at baseline, 4 samples on-treatment and 15 samples post-treatment) from MDACC and 20 samples (8 samples at baseline, 9 samples on-treatment and 3 samples post-treatment) from MGH were analyzed by ChIP-seq. We also analyzed 12 RNA-access samples (7 samples at baseline and 5 samples post-treatment) from MDACC and 32 RNA-seq samples (20 samples at baseline and 12 samples post-treatment) from MGH for RNA expression. Response rates were assessed based on RECIST criteria. Sex, age, disease stage, and progression time of each sample are included in **Table S1**.

### Cell lines

Short-term culture (STC) tumor cells and TILs pair were obtained from the same anti-PD-1– treated melanoma patient. Short-term culture tumor cells were cultured in RPMI and GlutaMAX supplemented with 10% fetal bovine serum, HEPES, human transferred insulin, and β- mercaptoethanol. TILs were cultured in RPMI and GlutaMAX supplemented with 10% human serum, sodium pyruvate, HEPES, human transferred insulin, β-mercaptoethanol and 3000IU/ml IL2 (PeproTech). The B16-F10 and BP melanoma cell lines and 293T cells were cultured in complete DMEM high-glucose medium, supplemented with 10% fetal bovine serum. All cell lines were cultured at 37°C with 5% CO2.

### Animal studies

All animal studies were performed according to the MDACC Institutional Animal Care and Use Committee (IACUC)–approved protocols. Five million B16-F10 or BP melanoma cells were injected subcutaneously into 6- to 8-week-old C57BL/6J mice (The Jackson Laboratory, #000664) and monitored every other day for tumor growth. On day 6 when tumors were palpable, mice with established tumor were randomly divided into 4 cohorts and treated every other day with IgG (100 μg/mouse), anti-PD-1 (100 μg/mouse), iBET-762 (7.5 mg/kg), or PBS (phosphate-buffered saline) via intraperitoneal injection until 14 days. Tumor volume was measured every other day. Mice were euthanized once any arm of the treatment developed tumors approaching or beyond the IACUC-approved limit of 1.5 cm.

### MDACC ChIP-Seq

ChIP was performed as described earlier (Terranova et al., 2018) with optimized shearing conditions and minor modifications. ChIP of 5-10 mg of flash-frozen patient melanoma tumors and mouse tumors were performed using 2 mg of antibody per ChIP experiment for H3K4me1 (#ab8895), H3K27ac (#ab4729), H3K4me3 (#ab8580), H3K79me2 (#ab3594), H3K4me9 (#ab8898), 3 mg of antibody per ChIP experiment for H3K27me3 (#ab6002), and 5 mg of antibody per ChIP experiment for BRD4 (#ab128874; all from Abcam). ChIP of 2-3 million of patient derived short-term culture tumor cells and TILs were performed using 5 mg of antibody per ChIP experiment for H3K27ac (#ab4729). Enriched DNA was quantified using Qubit (Thermo Fisher Scientific), and ChIP libraries were amplified and barcoded using the NEBNext Ultra II DNA library preparation kit (New England Biolabs) according to the manufacturer’s recommendations. Following library amplification, DNA fragments were size-selected (200-600 bp) using AMPure XP beads (Beckman Coulter), assessed using Bioanalyzer (Agilent Technologies), and sequenced at the Advanced Technology Genomics Core (MDACC) using Illumina HiSeq 2000 (36-bp single-end format).

### MGH ChIP-Seq

A total of 20-50 mg of snap-frozen melanoma tissues were pulverized by Geno/Grinder for 2 min at 1500 rpm and then fixed with 1% methanol-free formaldehyde plus protease inhibitor cocktails (Roche) for 10 min at room temperature and quenched by 125 μM glycine for 5 min at room temperature. Samples were incubated in cold radioimmunoprecipitation assay buffer (RIPA buffer: 50 mM Tris pH 8.0, 150 mM NaCl, 1% NP-40, 0.5% deoxycholate, 0.1% SDS) supplemented with protease inhibitor and sonicated using Covaris E220. Supernatants were quantified using Bio-Rad protein assay kits, and 1 mg of protein was loaded on 96-well plates for ChIP.

Protein A/G–coated silica columns embedded pipette tips were used for immunoprecipitating H3K27Ac antibody–bounded proteins instead of Protein A/G beads. The DNA was eluted in 100 μl 50 mM Tris pH 8.0 and 10 mM EDTA with 1% SDS after several washes, and the eluates were treated with proteinase K for 16 h at 65°C before library synthesis using NEBNext Ultra II DNA library preparation kits (New England Biolabs). The samples were sequenced on HiSeq 2000 (Illumina), and 30-50 million paired-end reads from each sample were recorded.

### ChIP-seq analysis

ChIP-seq data were quality controlled and processed by pyflow-ChIPseq (Tang, 2017a) a ChIP-seq pipeline based on snakemake (Koster and Rahmann, 2012). Briefly, raw reads were mapped by bowtie1 (Langmead et al., 2009) to the hg19 genome. Duplicated reads were removed, and only uniquely mapped reads were retained. RPKM-normalized bigwigs were generated by deep tools (Ramirez et al., 2016), and tracks were visualized with Integrative Genomics Viewer (Robinson et al., 2011). Narrow peaks were called using MACS1.4 (Zhang et al., 2008) with a p-value of 1e-5. For broad domains, the MACSv2.0.10 peak caller was used with a --broad-cutoff p-value of 1e-5. Chromatin state was called using ChromHMM (Ernst and Kellis, 2012), and the emission profile was plotted by ComplexHeatmap (Gu et al., 2016). Chromatin state models were learnt jointly on all data for all 6 histone marks (H3K4me1, H3K4me3, H3K27ac, H3K79me2, H3K9me3 and H3K27me3) from 25 melanoma tumors and a model with 15 states was chosen for detailed analysis. Heatmaps were generated using R package EnrichedHeatmap (Gu et al., 2018). Super-enhancers were identified using ROSE (Loven et al., 2013) based on H3K27ac ChIP-seq data.

### Chromatin State Transition Analysis

ChromHMM profiles of 6 pre-treatment non-responders and 5 pre-treatment responders were consolidated using epilogos. A pipeline was made to automate the calculation, and scripts used to re-code the ChromHMM states can be found at https://github.com/crazyhottommy/pyflow-chromForest/tree/vsurf_merge. With the output of epilogos, the chromatin state for each bin was chosen for the state that contained the greatest weights. A helper script can also be found at the link above. The consolidated ChromHMM profiles by epilogos were compared. The number of bins that switched chromatin states between groups was obtained. The number of bases that showed the transition change was obtained by multiplying the number of bins with the bin size (1000 bp). A Circos transition plot was made by the “circlize” R package. The script can be found in the GitLab repository https://gitlab.com/tangming2005/SKCM_IMT/blob/master/scripts/choose_state.py.

The consolidated ChromHMM profiles by epilogos were read into the R package EnrichedHeatmap. The chromatin state (categorical variable) was plotted in a 25-kb window centered on the active enhancer bins (chromatin state E7). Only bins that had E7 in one of the groups were retained for plotting. For 2-group comparisons, the bins were merged if the same change of state occurred in consecutive bins. A helper script can be found in https://gitlab.com/tangming2005/SKCM_IMT/blob/master/scripts/merge_bin.py.

### M-value processing and IDR calculations

To derive the M-values, we first used .bam files from both the ChIP and whole-cell extract files, along with a common peak file of 244,472 peaks, as inputs to MAnorm using default arguments. The common peak file was generated using the MACS2 “bdgdiff” function between combined pileups of responder and non-responder samples across the MDACC and MGH cohorts. The resulting normalized outputs from MAnorm were first used to filter samples by imposing an M > 0 and p < 0.05 filter. All samples had to have 20% of peaks bypassing the threshold, or they were discarded from the analysis. Thirty samples from the MGH cohort passed this filter, while 27 MDACC samples passed this filter. Next, we subjected the samples to the IDR algorithm. In this case, average M-values for all peaks were calculated for both cohorts, and the 2 average M-value vectors were utilized as inputs to the IDR algorithm with default arguments. The resulting 77,356 peaks were considered the final replicated peak set used for all downstream analyses.

### Differential H3K27ac ChIP activity calling

By leveraging the M-values, we determined the responder vs. non-responder differential response. We first batch-normalized the 2 cohorts’ M-values using the ComBat algorithm from the R package “sva” (Leek et al., 2012). Next, we used limma’s empirical Bayes modeling framework to construct a linear model regressing response and treatment time against M-values. We modeled patient identity—for patients with more than one sample analyzed— as a random effect. In order to be considered validated, a peak has to fulfill the nominal p-value cutoff of both the MDACC and MGH cohorts and the sign of the coefficients must be the same across the two cohorts. We note that this nominal p-value cutoff is ordinarily insufficient to control for false positive discoveries in a single cohort study. However, we require explicit confirmation for putative differential peaks in both MDACC and MGH cohort. The combined false positive rate for a gene to be falsely discovered in two distinct datasets is substantially lower than what the nominal p-value cutoff would suggest.

### Predicting immunotherapy response based on epigenomic features

We first stratified the dataset into 22 ChIP-seq and 26 RNA-seq cross-validation. Within each fold, the N=21 ChIP-seq and N=25 RNA-seq training examples were first stratified into MDACC and MGH cohorts. Within each cohort, we repeated the differential peak calling process to identify a set of replicated peaks at nominal p-value less than 0.05. We used the replicated peaks as features to a random forest binary classifier with 5 to 20 trees on the training data using the R package “randomForest”(Liaw and Wiener, 2002). We reported the ROC and auROC based off the classifier that had the highest LOO-CV auROC for each assay type. We then assessed the predictive of the performance by evaluating the predicted probability on the held out testing sample across all 22 ChIP-seq and 26 RNA-seq folds against the ground truth labels. We plotted the ROC and calculated the auROC using the R package “plotROC”(Sachs, 2017).

### Global test for groups of peaks

To run the global test for genes, we first associated each of the peaks in the common peak set with a gene via the HOMER (Heinz et al., 2010) annotatePeak function. Each gene’s associated peaks were organized as a group for the global test. The global test was conducted using the function “gt” with default parameters using the “globaltest” R package (Goeman et al., 2004).

### RNA Access sequencing and analysis of MDACC tumors

mRNA libraries of the melanoma tumor (n = 12) samples were prepared from 200 ng of total RNA using the TruSeq Stranded mRNA HT Sample Preparation Kit. Samples were dual-indexed before pooling. Libraries were quantified by qPCR using the NGS Library Quantification Kit. Pooled libraries were sequenced using the HiSeq 2000 (Illumina) according to the manufacturer’s instructions. An average of approximately 30 million paired-end reads per sample were obtained.

The quality of raw reads was assessed by using FastQC (https://www.bioinformatics.babraham.ac.uk/projects/fastqc/). The raw reads were aligned to the Homo sapiens genome (hg19) using STAR 2.4.2a (Dobin et al., 2013) (https://github.com/alexdobin/STAR/releases/tag/STAR_2.4.2a). The mappability of unique reads on average was ∼89% RNA-seq dataset. The raw counts were computed using the quantMode function in STAR. The read counts that were obtained are analogous to the expression level of each gene across all the samples. Genes with raw mean reads of greater than 10 were used for normalization and differential gene expression analysis using the DESeq2 (Love et al., 2014) package in R. Genes with an absolute log2 fold-change greater than log2(1.5) and p < 0.05 were called as differentially expressed genes. SKCM TCGA RNA-seq transcription comparison analysis was performed on the UALCAN website (Chandrashekar et al., 2017).

### RNA-seq and analysis of MGH tumors

Total RNA from 5-20 mg of melanoma primary and metastatic tissues was extracted using AllPrep DNA/RNA Mini isolation kit (Qiagen). A total of 100 ng of total RNA was used as input for RNA-seq libraries using SMARTer Stranded Total RNA-seq - Pico input (Takara Bio USA, Inc.) to remove rRNA transcripts. Each library was sequenced on HiSeq 2000 (Illumina), and approximately 20 million single-ended reads were recorded. Reads were aligned to *Homo sapiens* reference hg38 using STAR 2.5.3. Read counts were quantified using featureCounts. Differential expression was performed via limma-voom (Ritchie et al., 2015). Multiple biological replicates stemming from the same patient were treated as a random effect, whereas batch effects were treated as a fixed effect.

### RNA-Seq analysis of murine tumor cells

mRNA libraries of the mouse melanoma tumor (n = 12) samples were prepared and sequenced using the HiSeq 2000 (Illumina). RNAseq data were processed by pyflow-RNAseq (Tang, 2017b), a snakemake based RNAseq pipeline. Raw reads were mapped by STAR (Dobin et al., 2013), RPKM normalized bigwigs were generated by *deeptools* (Ramirez et al., 2016), and gene counts were obtained by *featureCount* (Liao et al., 2014). Differential expression analysis was carried out using *DESeq2* (Love et al., 2014). Gene set enrichment analysis (GSEA) was done using the GSEA tool (Subramanian et al., 2005) in pre-rank mode. The signed fold change *–log10(pvalue) metric was used to pre-rank the genes.

### WGS data analysis and TMB calculation

Whole genome sequencing data from 34 anti PD-1 treated melanoma patient samples were aligned to human reference genome version hg38 using the Burrows-Wheeler Alignment tool (v.0.7.17), and duplicate removed by samtools (v.1.15). Somatic single nucleotide variations (SNVs) were identified using Mutect 2 (v.4.2.4.1), and variants likely to be germline were filtered out by gnomAD (v.2) and FilterMutectCalls. Tumor mutation burden was defined as the number of non-synonymous mutations in the coding region per megabase.

### HiChIP and data analysis

HiChIP experiments were performed as previously described by Mumbach et al. (Mumbach et al., 2016), with minor modifications. Briefly,1 × 10^7^ ICB resistant STC cells were crosslinked. In situ contacts were generated in isolated and pelleted nuclei by DNA digestion with MboI restriction enzyme, followed by biotinylation of digested DNA fragments with biotin–dATP, dCTP, dGTP, and dTTP. Thereafter, DNA was sheared with Covaris E220 with the following parameters: fill level = 10, duty cycle = 5, PIP = 140, cycles/burst = 200, and time = 4 min; ChIP was done for H3K27Ac using the anti-H3K27ac antibody. After reverse-crosslinking, 150 ng of eluted DNA was taken for biotin capture with Streptavidin C1 beads followed by transposition with Tn5. In addition, transposed DNA was used for library preparation with Nextera Ad1_noMX, Nextera Ad2.X primers, and Phusion HF 2X PCR Master Mix. The following PCR program was performed: 72°C for 5 mins, 98°C for 1 min, then cycle at 98°C for 15 s, 63°C for 30 s, and 70°C for 1 min. Afterward, libraries underwent double-sided size selection with AMPure XP beads. Finally, libraries were paired-end sequenced with reading lengths of 76 nucleotides. HiChIP paired-end reads were aligned to the MboI-digested hg19 genome using the HiC-Pro pipeline with default conditions. The default setting of HiC-Pro removes duplicate reads, assigns reads to MboI fragments, identifies valid interactions, and generates high-resolution interaction matrices. HiChIP for H3K27ac generated high-resolution contact maps containing ∼65 million valid interactions in STC2765 cells. Files for Juicebox visualization were generated using the HiC-Pro hicpro2juicebox.sh command based on the total valid interactions. H3K27ac-mediated loops were identified with the hichipper/diffloop programs using the HiC-Pro (Servant et al., 2015) output and ChIP-seq peaks from H3K27ac as anchor loci. *Hichipper* identifies intrachromosomal looping between anchor loci within 5 kb to 2 MB and produces a per-loop FDR value from the loop proximity bias correction implemented by Mango. Using the Mango output from hichipper (Lareau and Aryee, 2018), diffloop was used to filter significant loops (FDR < 0.01, width ≥ 5000, loop-count ≥ 2) and define enhancer-enhancer and enhancer-promoter interactions.

### Enhancer data analysis – peak-to-gene linking predictions

To identify putative causal links between enhancer peaks and gene expression, we used a HiChIP based approach. Enhancer-promoter interaction catalogs from STC2765 Hi-ChIP data and from a previous publication (Cao et al., 2017) was overlapped with the query enhancer peaks in order to obtain its taget refseq promoter. In addition, we also used the ChIPseeker package for annotation, using addFlankGeneInfo function for SEs.

### Pathway analysis

Differential enhancer-associated genes in each group were imported into the clusterProfiler (Yu et al., 2012) or Consensus PathDB (http://cpdb.molgen.mpg.de/) for pathway analysis, restricted to Gene Ontology, KEGG, Hallmark, and WiKiPathways gene sets. The “enrichplot” package (Yu, 2019) was used to generate dot plots and networks for gene sets enriched with an FDR cut-off of < 0.05.

### Enrichment of motifs in cell-specific enhancer peaks

To identify the motifs over-represented within each enhancer peak sets, we used the HOMER motif database and the coordinates of melanoma cells or TILs specific peak sets.

### Enhancer modulation using CRISPR-dCas9-KRAB

To modulate gene expression without altering the target DNA sequences, an RNA-guided, catalytically inactive Cas9 (dCas9) fused to a transcriptional repressor domain (KRAB) was used to silence genomic regions identified as enhancers via KRAB repression at the promoter region. To generate a dCas9-KRAB effector stable cell line, we produced lentiviral particles from pHAGE EF1α-dCas9-KRAB (Addgene plasmid #50919) using Pax2 and VSVg. Transduced cells were selected for 6 days with the use of antibiotic resistance and were expanded to generate a stable cell line.

Next, gRNAs were designed by using the GPP Web Portal of the Broad Institute. gRNAs sequences are listed in **Table S6**. Annealed gRNA oligos were ligated to pLKO.1-puro U6 sgRNA BfuAI stuffer (Addgene plasmid #50920), and lentiviral particles were generated. A transduction procedure was performed in the stable dCas9-KRAB cell line, and transduced cells having both dCas9-KRAB and gRNA constructs were selected with the use of antibiotic resistance. To evaluate the effects of the recruitment of dCas9-KRAB to the target enhancer’s genomic region, H3K27ac ChIP followed by quantitative PCR for enhancer regions was performed to assess the enrichment level of H3K27ac at the enhancer site in modulated cells compared with the non-modulated parental control cells. To investigate the impact of enhancers’ modulation on the corresponding gene expression, qRT-PCR was performed for the target gene.

### RT-qPCR

RNA was isolated using RNeasy kit (qiagen) using manufacturer’s protocol. cDNA was prepared using SuperScript III first strand synthesis kit (Thermo Fisher) using 2ug of RNA and manufacturer’s protocol. Quantitative PCR was performed using QuantiTect Sybr Green PCR kit in Stratagene’s Mx3000p system.

### *In vitro* inhibitor assays

Melanoma short-term culture line STC2765 were treated with crizotinib (2 μM, 24 h) or iBET-762 (1 μM, 72 h) prior to co-culture with TIL2765 cells.

### TILs and matched tumor cells co-culture

Tumor cells were labeled with DDAO-SE followed by addition of an effector cell suspension to achieve the desired effector:target ratio. These co-cultures were incubated at 37°C in 5% CO2 in a humidified incubator for 3 h. The cells were fixed and permeabilized with Cytofix/Cytoperm solution (BD Biosciences, #554722) for 20 min at RT immediately. The cells were stained for 30 min on ice with 5 μl of biotin-labeled anti–cleaved caspase-3 monoclonal antibody (BD Biosciences, #550821). The cells were washed in Perm/Wash buffer (BD Biosciences, #554723) 2 times and re-suspended in PBS and 1% fetal bovine serum for analysis on a flow cytometer.

### Flow cytometry

TILs were stained with fluorochrome-conjugated monoclonal antibodies (CD3, CD4, and CD8 from BD Biosciences) in FACS wash Buffer (Dulbecco’s phosphate buffered saline 1× with 1% bovine serum albumin) for 30 min on ice for surface staining. Dead cells were excluded using Ghost Dye^TM^ Violet 450 cell viability dye from Tonbo Biosciences. For intracellular staining of active caspase-3, cells were fixed and permeabilized using Cytofix/Cytoperm (BD Biosciences) and stained with anti–cleaved caspase-3 (BD Biosciences) on ice as well. Stained cells were acquired using BD FACSCanto II and analyzed using FlowJo software (Tree Star).

### Survival analysis

The “survminer” package was used for drawing the Kaplan-Meier plots and defining the optimal threshold (function surv). The outcome was overall survival censored at 10 years. p-values reported for the univariate model corresponded to the log-rank test.

### Statistical analysis

The 2-tailed Student *t*-test was used to determine the statistical significance of 2 groups of data using GraphPad Prism. Data are presented as means ± standard error of the mean (SEM, error bars) of at least 3 independent experiments or 3 biological replicates. p-values less than 0.05 were considered statistically significant. *, p < 0.05; **, p < 0.01; and ***, p < 0.001. Correlation of expression level between BRD4 and CD274(PD-L1) was computed with nonparametric Spearman’s rank correlation coefficient.

### Data and code availability

All ChIP-Seq dataset generated from ICB treated melanoma tumors have been deposited into the Gene Expression Omnibus (GEO) repository (accession #GSE171283). All codes are available at https://gitlab.com/railab.

## SUPPLEMENTARY FIGURE LEGENDS

**Figure S1:**
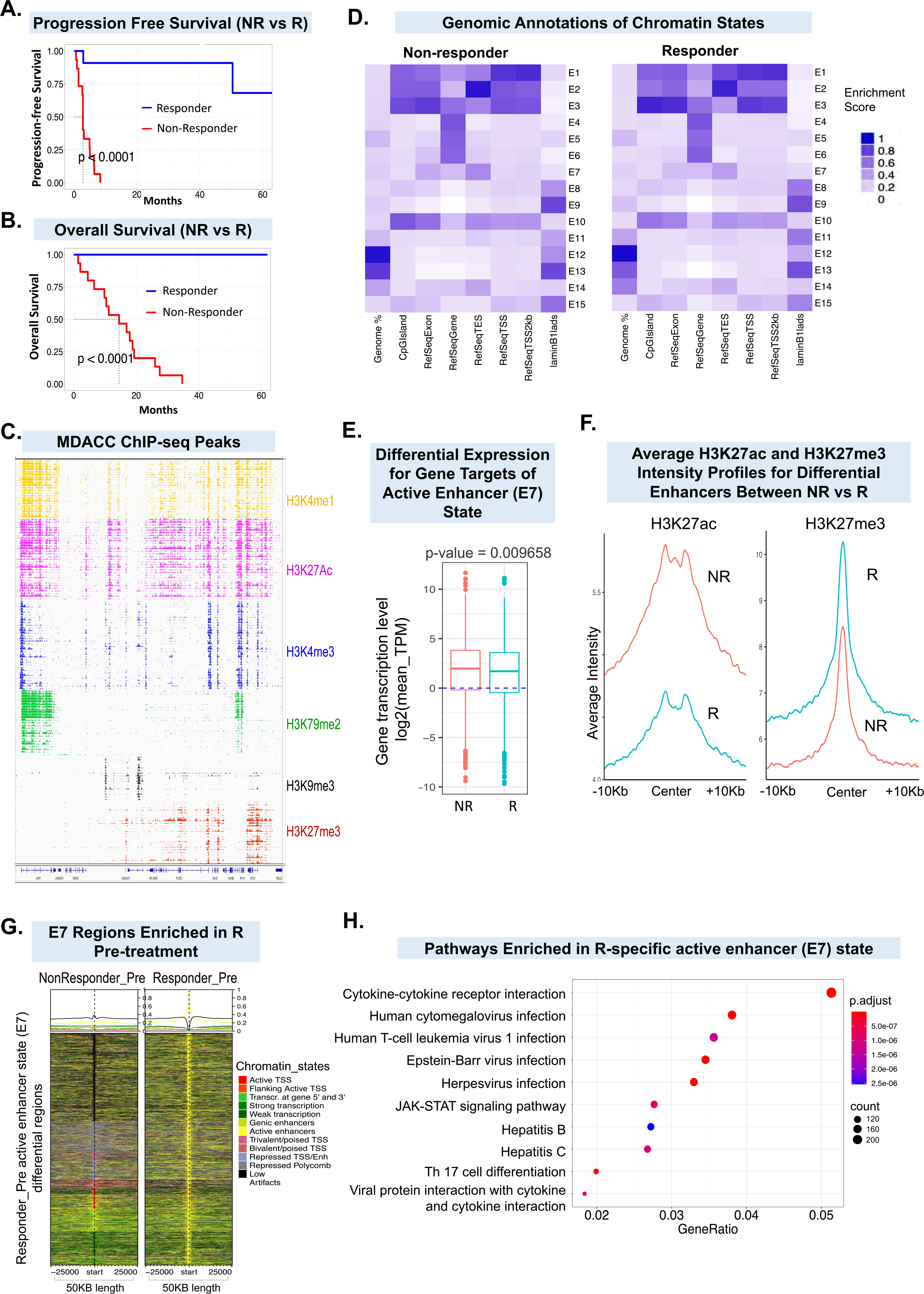
Chromatin state differences between anti-PD-1 responders and non-responders. **A-B.** Kaplan-miere curve showing progression-free survival (**A**) and overall survival (**B**) for responders and non-responders to anti-PD-1 therapy in melanoma (MDACC cohort). The log-rank (Mantel-Cox) p value is shown for the difference in survival. **C.** IGV view of 6 different histone mark profiles (as noted on the right side) on the shown chromosomal region in all the anti-PD-1–treated patients. **D.** Genomic annotation enrichments for each chromatin state in anti-PD-1 non-responder (top) and responder (bottom) tumor samples. **E.** Box plots showing the log2 mean expression levels (Transcripts Per Million, TPM) of genes associated with enhancer state E7. Genes were linked using H3K27ac HiChIP data from STC2765 melanoma cell lines and FANTOM data as described in the methods section. The bottom and the top of the rectangles indicate the first quartile (Q1) and third quartile (Q3), respectively. The horizontal lines in the middle signify the median (Q2), and the vertical lines that extend from the top and the bottom of the plot indicate the maximum and minimum values, respectively. **F.** Average intensity plots for H3K27ac (left) and H3K27me3 (right) on loci that lost H3K27ac marks (from Fig. 1E) in pre-treatment R versus NR tumors from MDACC cohort. **G.** Heatmap of chromatin state intensities for 20,194 loci that showed a switch from E7 (yellow) in responder pre-treatment samples (right) to any other state in non-responder pre-treatment samples (left), as shown by colors for each state. **H.** Dot plot showing the pathways in genes targeted by E7 state enhancers that were significantly enriched in responders compared to non-responders. Dot size represents the gene counts; adjusted p-values are shown and are color-coded based on the level of significance.

**Figure S2:**
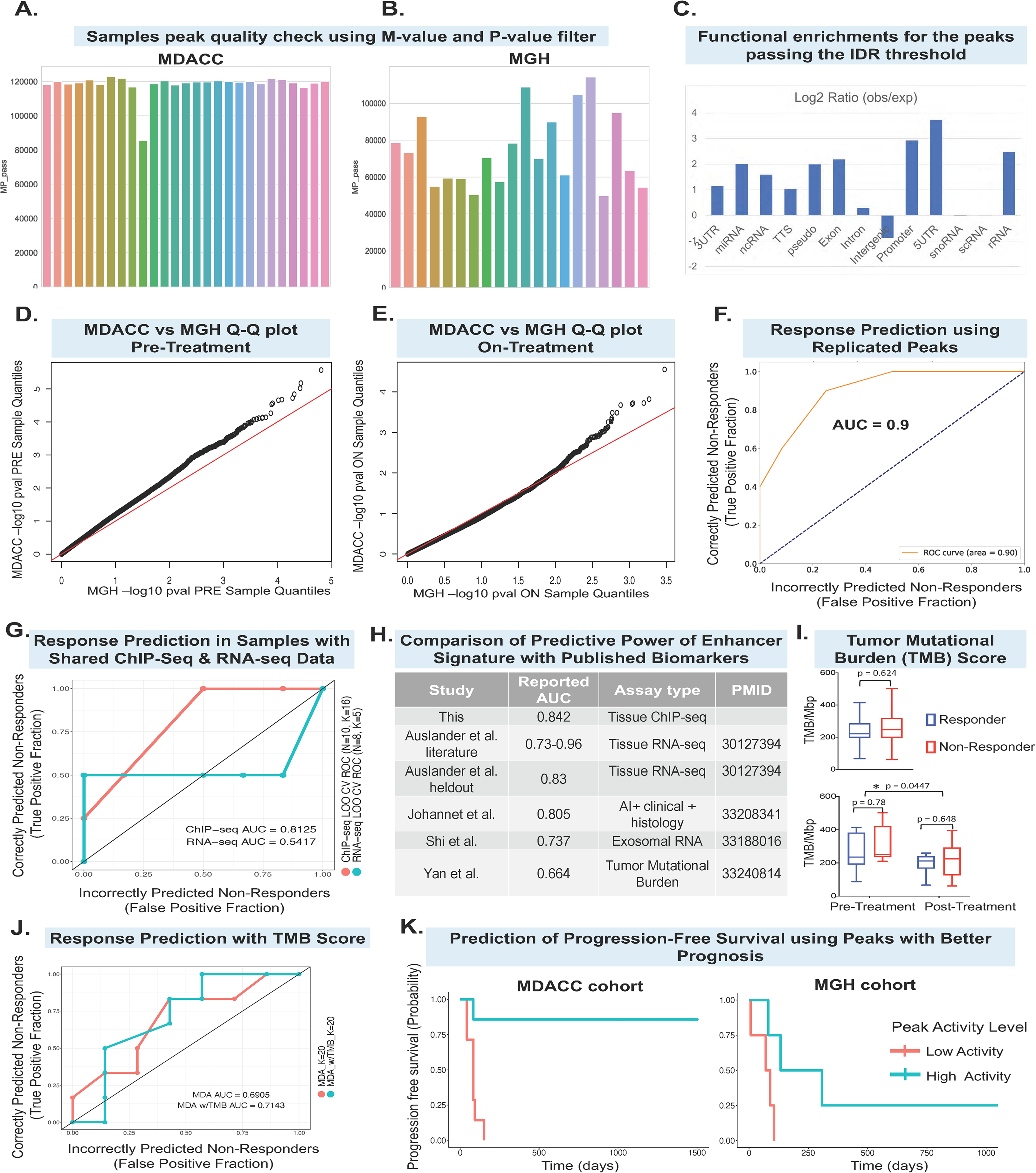
Validation of enhancer signature between MDA and MGH cohorts. **A.** Bar chart showing number of peaks in individual samples from MDACC cohort that pass quality threshold of M-value>0 and MAnorm p>0.1. **B.** Bar chart showing number of peaks in individual samples from MGH cohort that pass quality threshold of M-value>0 and MAnorm p>0.1. **C.** Functional enrichments for the 84,317 peaks passing the IDR threshold. **D.** QQ-plot between MGH (x-axis) and MDACC (y-axis) sample quantiles from the pre-treatment (PRE) comparison. **E.** QQ-plot between MGH (x-axis) and MDACC (y-axis) sample quantiles from the on-treatment (ON) comparison. **F.** Receiver operating characteristic (ROC) of random forest trained predictive models utilizing the 437 replicated pre-treatment peaks. The ROC curve was formed by concatenating predictions from 2 models: a model trained exclusively on MGH data and tested on MDACC data, and a model trained exclusively on MDACC data and tested on MGH data. **G.** Receiver operating characteristic (ROC) of random forest trained predictive models using replicated ChIP-seq peaks or RNA-seq genes across N=22 pre-treatment ChIP-seq samples and N=26 pre-treatment RNA-seq samples. The features within each cross-validation fold were determined by finding the set of replicated peaks or genes across the MDACC and MGH cohorts by computing the set of intersecting peaks or genes that were nominally significant in both the MDACC & MGH cohorts in the training cohort. We analyzed a total of 23,457 RNA-seq genes and 84,317 ChIP-seq peaks. This was used to train a random forest within each training set with K=5 to K=20 trees, the reported the ROC & auROCs are derived from the best performing random forest classifier. The ROC curve and auROC was formed by concatenating predictions from the N=10 ChIP-seq and N=8 RNA-seq shared samples (samples with both RNA-seq and ChIP-seq from the pre-treatment timepoint) across cross validation folds. **H.** Comparison of observed LOO CV auROC and literature auROC across melanoma checkpoint blockade response prediction studies. **I.** Box plots showing Tumor Mutational Burden (TMB) in all responder vs non-responder patients (top), in pre- or post-treatment responder vs non-responder patients (bottom). The bottom and the top of the rectangles indicate the first quartile (Q1) and third quartile (Q3), respectively. The horizontal lines in the middle signify the median (Q2), and the vertical lines that extend from the top and the bottom of the plot indicate the maximum and minimum values, respectively. **J.** Effect of TMB as a predictive feature for pre-treatment response prediction in the MDACC cohort. Here we evaluate two alternative models for predicting pre-treatment outcomes using ChIP-seq data in the MDACC cohort. In LOO-CV across N=13 MDACC samples with both ChIP-seq and TMB data, we observed incorporating TMB data along with differential ChIP-seq peaks (AUC=0.7143) as features to a random forest classifier with K=20 trees resulted in a slightly increased AUC compared to only using differential ChIP-seq peaks alone (AUC=0.6905). **K.** Kaplan-Meier plots showing progression-free survival in MDACC (left) or MGH (right) cohorts for 3 out of the 32 peaks which offered better prognosis as a result of increased peak signal. The normalized ChIP activity values were studentized across the MDACC cohort, then the median value was used to determine the high-activity vs. low-activity groups. There was a total of 8 patients (4 low enhancer activity, 4 high enhancer activity) in the MGH group and a total of 14 patients (7 low enhancer activity, 7 high enhancer activity) in the MDACC group. The peaks were selected using a p=0.05 cutoff for the Cox proportional hazards test.

**Figure S3:**
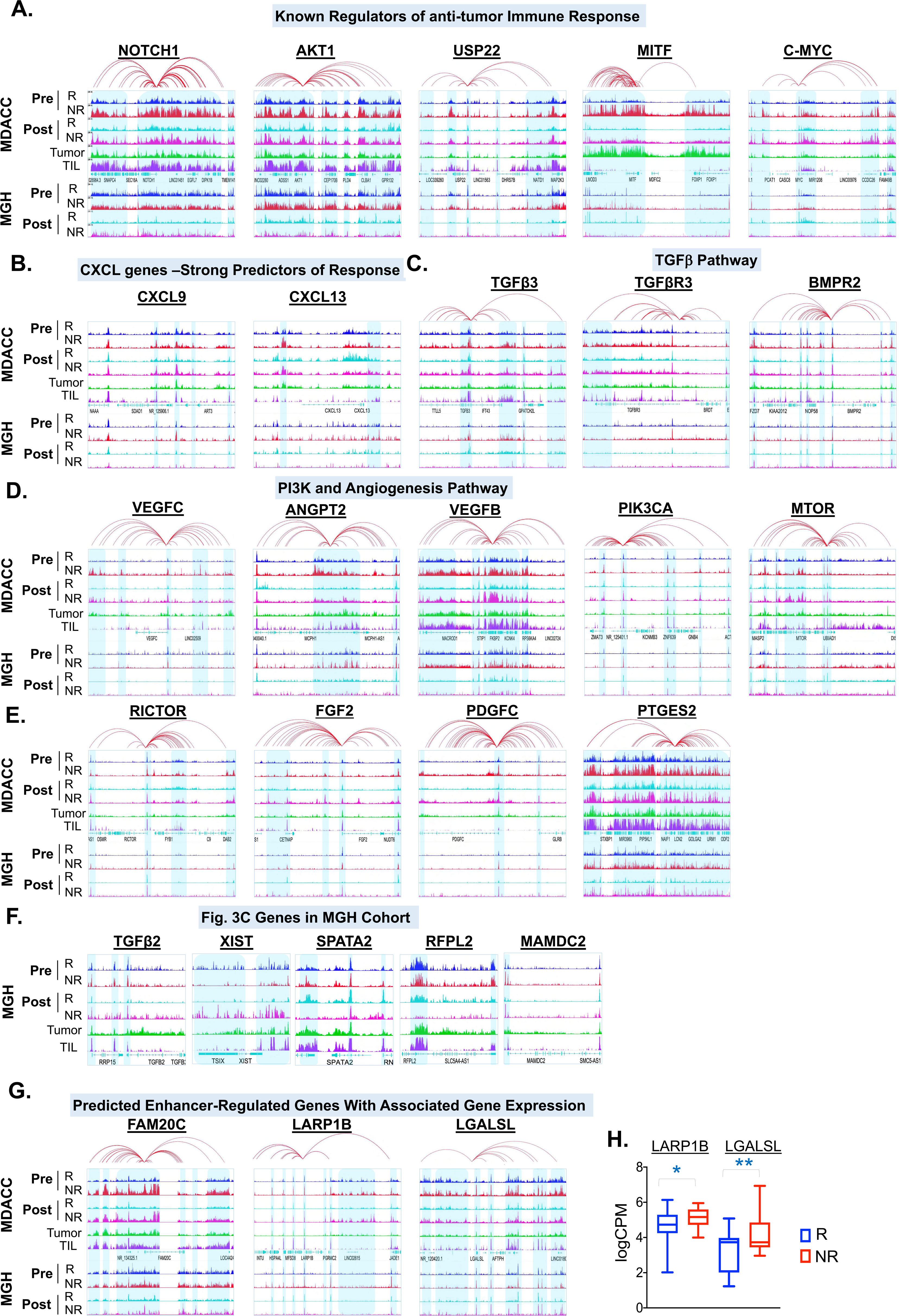
Differences in enhancer activation on specific groups of genes between non-responders and responders to anti-PD-1. **A.** IGV snapshot of aggregate H3K27ac profiles around NOTCH1, AKT1, USP22, MITF and c-MYC in NR and R samples from both cohorts as well as isolated melanoma STCs or TILs. The red line loops in all panels in this figure depict E-P interactions identified from H3K27ac HiChIP data from STC2765 cells and/or previously predicted E-P networks (Cao et al., 2017). **B.** IGV snapshot of aggregate H3K27ac profiles around CXCL9 and CXCL13 in NR and R samples from both cohorts as well as isolated melanoma STCs or TILs. **C-E.** IGV snapshot of aggregate H3K27ac profiles around TGFβ3, TGFβR3, BMPR2, VEGFC, ANGPT2, VEGFB, PIK3CA, MTOR, RICTOR, FGF2, PDGFC and PTGES2 in NR and R samples from both cohorts as well as isolated melanoma STCs or TILs. F. IGV snapshot of aggregate H3K27ac profiles around TGFβ2, XIST, SPATA2, RFPL2, and MAMDC2 in MGH cohort NR samples, R samples, isolated melanoma STCs, or isolated TILs. G. IGV snapshot of aggregate H3K27ac profiles around FAM20C, LARP1B and LGALSL in NR and R samples from both cohorts as well as isolated melanoma STCs or TILs. H. Volcano plot showing MDACC and MGH cohort combined expression data differentially expressed genes (blue and red) in responder vs non-responders. X-axis shows log fold change (FC), and y-axis represents p-value of gene expression change. I. Box plot showing the gene expression level of LGALSL and LARP1B genes in non-responder and responder pre-treatment samples. In the box plot, the bottom and the top of the rectangles indicate the first quartile (Q1) and third quartile (Q3), respectively. The horizontal lines in the middle signify the median (Q2), and the vertical lines that extend from the top and the bottom of the plot indicate the maximum and minimum values, respectively.

**Figure S4:**
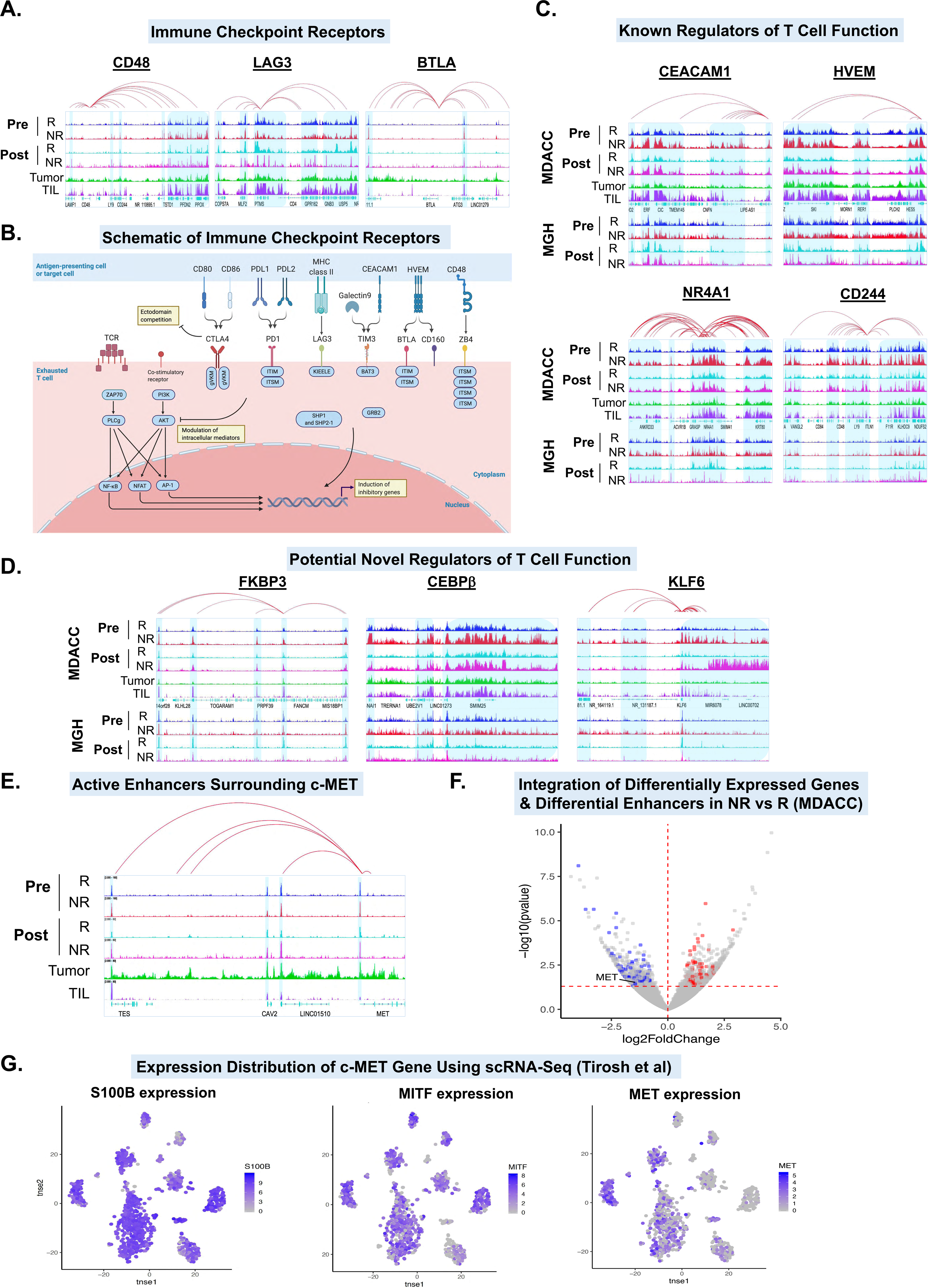
Gene targets of activated enhancers in anti-PD-1 non-responders. **A.** IGV snapshot of aggregate H3K27ac profiles around CD48, LAG-3, and BTLA in NR and R samples from both cohorts as well as isolated melanoma STCs or TILs. The red line loops in all panels in this figure depict E-P interactions identified from H3K27ac HiChIP data from STC2765 cells and/or previously predicted E-P networks (Cao et al., 2017). **B.** Schematic showing the key immune checkpoint receptors on exhausted TILs. **C-D.** IGV snapshot of aggregate H3K27ac profiles around CEACAM1, HVEM, NR4A1 and CD244 (**C**); and FKBP3, CEBPB, and KLF6 (**D**) in NR and R samples from both cohorts as well as isolated melanoma STCs or TILs. E. IGV snapshot of aggregate H3K27ac profiles around c-MET in NR and R samples from MGH cohort as well as isolated melanoma STCs or TILs. F. Volcano plot showing MDACC cohort differentially expressed genes (gray dots) and differentially enriched enhancers targeted genes (red or blue) in R vs. NR samples. X-axis shows log2 fold change, and y-axis represents p-value of gene expression change. G. Distribution of expression of S100B, MITF, and MET genes in 2-dimensional embedding obtained by tSNE. Data were extracted from melanoma single-cell RNA-seq data (Tirosh et al., 2016). Each cell is colored according to the gene expression level.

**Figure S5:**
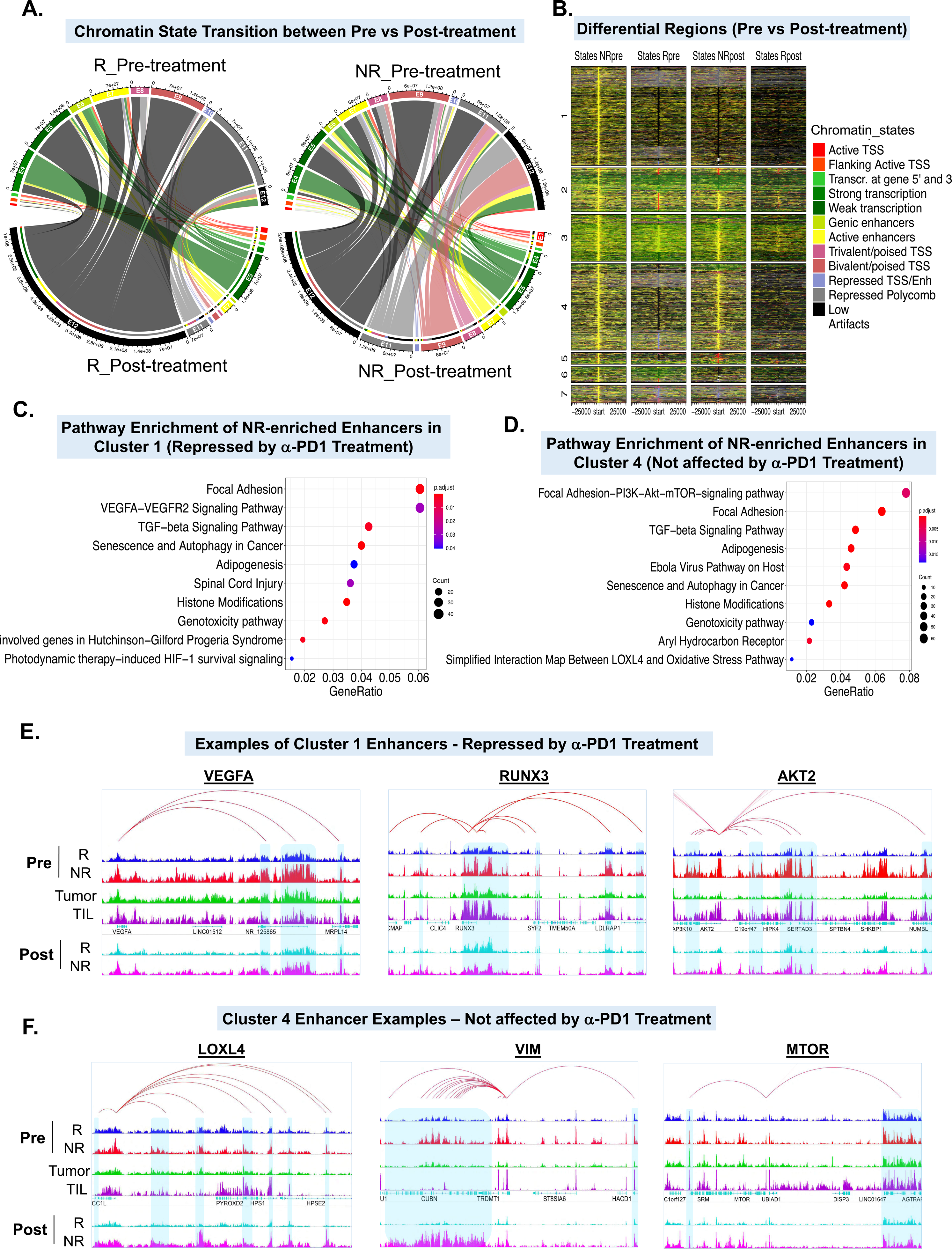
Chromatin state transitions during non-response to immunotherapy. **A.** Circos plot showing chromatin state switches between pre-treatment and post-treatment samples from responders (left) or non-responders (right). Chromatin state transitions were calculated based on epilogos (see Methods). Yellow color bands show high percentage of active enhancer state E7 transitioning to low or repressed states E12 or E11. **B.** Heatmap of chromatin state intensities for 31,155 loci that show switch from E7 in NR pre-treatment samples to any other state in R pre-treatment, NR post-treatment, or R post-treatment samples as shown by colors for each state. Y-axis shows the clusters (numbered one through seven) of genomic loci that follow specific transition patterns. **C-D.** Dot plot representation of significantly enriched pathways in gene targets of enhancers present in Cluster 1 (**C**) and Cluster 4 (**D**) from panel **B**. Dot size represents the gene counts. Adjusted p-values are color-coded based on the level of significance. **E-F.** IGV snapshot of aggregate H3K27ac profiles around VEGFA, RUNX3, and AKT2 (**E**) and LOXL4, VIM, and MTOR (**F**) in NR and R samples from both cohorts as well as isolated melanoma STCs or TILs. Highlighted regions depict specific enrichment of H3K27Ac enhancer peaks in NR pre-treatment (**E**) or post-treatment (**F**) samples. The red line loops in all panels in this figure depict E-P interactions identified from H3K27ac HiChIP data from STC2765 cells and/or previously predicted E-P networks (Cao et al., 2017).

**Figure S6:**
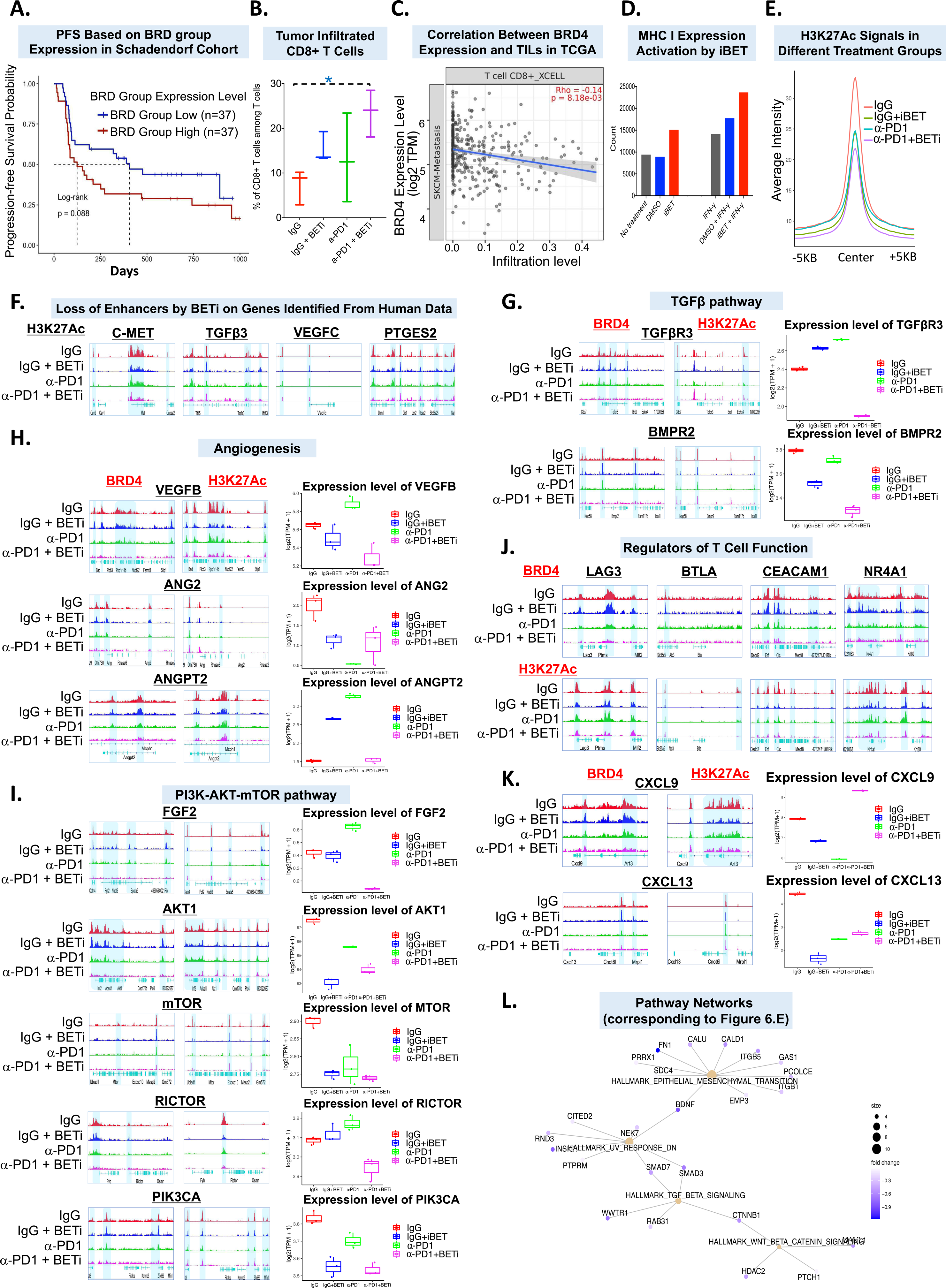
Molecular mechanism behind combination treatment of bromodomain inhibitors plus anti-PD-1. **A.** Kaplan-miere curve showing progression-free survival in two groups of patients: one bearing high levels of BRD2, BRD3 and BRD4 expression, and second bearing low levels of these BRD proteins in Schadendorf cohort of anti-PD-1 treated patients. **B.** Graph showing the flow cytometry analysis results of infiltrated CD8+ T-cell percentages in tumors derived from experiment shown in Fig.5F. **C.** Scatter plot showing the correlation between BRD4 expression and TIL infiltration in the TCGA melanoma cohort. **D.** Bar graph showing MHC class I expression in STC2765 cells which were untreated or treated with DMSO or iBET-762 (1μM, 72 hours) alone or along with IFN-γ. **E.** Average intensity curves of ChIP-Seq reads (RPKM) for H3K27ac in tumors treated with IgG, α-PD-1, IgG + iBET-762 or α-PD-1 + iBET-762 (corresponding to experiment shown in Fig. 5E) at all enhancer regions. Enhancers are shown in a 10kb window centered on the middle of the locus. **F.** IGV snapshot of H3K27Ac ChIP-seq signal around c-MET, TGFβ3, VEGFC and PTGES2 gene loci in tumors treated with IgG, α-PD-1, IgG + iBET-762 or α-PD-1 + iBET-762. **G-I.** IGV snapshot of aggregate BRD4 and H3K27ac (left) profiles around TGFβR3, BMPR2, VEGFB, ANG2, ANGPT2, FGF2, AKT1, MTOR, RICTOR and PIK3CA in in tumors treated with IgG, α-PD-1, IgG + iBET-762 or α-PD-1 + iBET-762. Box plot (right) shows mRNA expression of genes shown in panels **F-H** (left). Each dot represents a sample. Colors represent 4 treatment groups shown in the plot. In the box plot, the bottom and the top of the rectangles indicate the first quartile (Q1) and third quartile (Q3), respectively. The horizontal lines in the middle signify the median (Q2), and the vertical lines that extend from the top and the bottom of the plot indicate the maximum and minimum values, respectively. J. IGV snapshot of aggregate BRD4 (top) and H3K27ac (bottom) profiles around LAG3, BTLA, CEACAM1 and NR4A1 in in tumors treated with IgG, α-PD-1, IgG + iBET-762 or α-PD-1 + iBET- 762. K. Left, IGV snapshot of aggregate BRD4 (top) and H3K27ac (bottom) profiles around CXCL9 and CXCL13 in tumors treated with IgG, α-PD-1, IgG + iBET-762 or α-PD-1 + iBET-762. Right, box plot representation of the mRNA expression level of CXCL9 and CXCL13. Each dot represents a sample. Colors represent 4 treatment groups shown in the plot. In the box plot, the bottom and the top of the rectangles indicate the first quartile (Q1) and third quartile (Q3), respectively. The horizontal lines in the middle signify the median (Q2), and the vertical lines that extend from the top and the bottom of the plot indicate the maximum and minimum values, respectively. L. Pathway network analysis for 107 genes from Fig. 6D-E obtained from overlap of human and mouse data.

## SUPPLEMENTARY TABLES

Table S1. Details of the patient samples utilized in this study and quality matrix of the generated chromatin data.

Table S2. List of significantly differentially enriched peaks and differentially expressed genes between responder and non-responder pre-treatment samples in MDACC and MGH cohorts as well as replicated peaks with annotation.

Table S3. List of H3K27ac peaks that are derived from overlap of replicated NR-specific and R-specific enhancers with tumor-specific or TIL-specific enhancers.

Table S4. List of active enhancer regions in cluster 1 and cluster 4 derived during analysis of chromatin state transitions between pre-treatment to post-treatment samples.

Table S5. List of genomic regions and associated genes that display loss of BRD4 binding and reduced expression in tumors treated with iBET-762 plus anti-PD-1 versus IgG.

Table S6. List of gRNAs used in the enhancer editing experiment shown in Figure 4.

